# Natural product-derived azaphilones are selective scaffolds for covalent lysine liganding in live cells

**DOI:** 10.64898/2026.09.27.751175

**Authors:** Wei Ding, Sophie Brameyer, Susanne H. Kirsch, Birthe Sandargo, Frank Surup, Stephan M. Hacker, Kirsten Jung, Stephan A. Sieber

## Abstract

Lysine-directed covalent liganding offers a promising strategy to expand the covalently ligandable proteome beyond cysteine. However, broadly applicable and selective lysine-reactive scaffolds remain limited. Here, we designed SCL-alkyne, a simplified azaphilone probe derived from the natural product sclerotiorin (SCL), for the global profiling of lysine reactivity in bacteria. SCL-alkyne displayed high intrinsic selectivity for lysine *in vitro* and in live *Escherichia coli* cells. *In situ* proteome profiling revealed that 84% of engaged sites belong to lysine residues and the protein *N*-terminus. Reactivity mapping provided an inventory of highly reactive sites showing that only a limited subset of reactive lysines is located in functional regions of proteins. Analysis of proteins enriched *via* the SCL-alkyne probe unraveled bacterial chemotaxis as a major target pathway which was confirmed by an impaired chemotactic response upon treatment of bacteria with SCL-alkyne. Competitive profiling of three structurally related azaphilone natural products identified 60 protein targets belonging to pathways related to iron-sulfur cluster assembly and antioxidant function. This study not only highlights the diverse modes-of-action of natural product azaphilones, but also establishes its core scaffold as effective lysine-reactive chemotype for identifying ligandable lysines.

## Introduction

Natural products provide a rich source of electrophilic warheads capable of covalently engaging diverse protein targets.^[1-2]^ These include *Michael* acceptors, nitriles, aldehydes, and strained ring systems such as epoxides and β-lactams.^[1-5]^ Although natural product-based covalent drugs have been on the market for decades, their importance for *de novo* drug design has only recently been recognized with the advent of targeted covalent inhibitors (TCIs) and a broader appreciation of their key advantages over reversible binders.^[6-8]^ Irreversible target engagement, for instance, can prolong pharmacological activity over the lifetime of the protein, largely independent of binding kinetics.^[8-9]^ The value of this concept has been successfully demonstrated by the approval of afatinib and ibrutinib, kinase inhibitors that covalently target cysteine residues near the active site *via* a strategically positioned *Michael* acceptor.^[10-11]^ In addition, screening of chemical libraries bearing electrophilic warheads has led to the discovery of unprecedented binding sites in proteins previously believed to be undruggable.^[12-13]^ Covalent liganding has, therefore, emerged as a powerful strategy, driven by rapid advances in fragment design, including approaches that address stereochemically defined pockets, by the expansion of chemistries beyond cysteine targeting, and by analytical methods that enable binding sites to be identified either through direct labeling or by competition-based activity-based protein profiling (ABPP).^[14-18]^

So far, the vast majority of covalent liganding approaches has been dedicated to the targeting of cysteine residues, in part due to the well-established chemistry of selective warheads such as *Michael* acceptors.^[19-20]^ However, cysteine accounts for only 1.7% of all amino acid residues present in proteins and is, therefore, absent from many desired binding sites.^[21]^ To increase overall coverage and enhance flexibility, other nucleophilic amino acids such as lysine, glutamic acid/aspartic acid, and tyrosine have come into focus, along with the development of tailored electrophiles.^[21-23]^ In particular, lysine, which accounts for 6% of amino acid residues in proteins, has attracted attention as a prevalent residue with established liganding approaches.^[24-25]^ Lysine-targeting strategies can be broadly divided into several classes based on their structural features: activated esters/amides, benzaldehydes, sulfur(VI) electrophiles and others (Figure 1A).^[24,26-33]^ These strategies have been successfully applied to covalent ligand discovery in both human and bacterial proteomes.^[34]^ For example, **STP-alkyne** enabled the labeling of 2,943 lysine sites across 804 proteins in *Staphylococcus aureus* SH1000, demonstrating broad proteome coverage.^[18]^ In addition, lysine targeting can provide an alternative strategy, when cysteine-based targeting is compromised by mutation. For instance, diethenyl sulfoximine (DESI) modifies lysine through a cyclization mechanism, in which the ε- amino group undergoes double addition to two vinyl groups to form a six-membered cyclic adduct.^[33]^ This approach has been used to develop a covalent allosteric EGFR inhibitor capable of overcoming the C797S mutation.^[33]^ However, the high intrinsic reactivity of some strategies could lead to low amino acid selectivity and poor biocompatibility, including cytotoxicity, thereby limiting their applications in live cells.^[27,29]^

**Figure 1.**
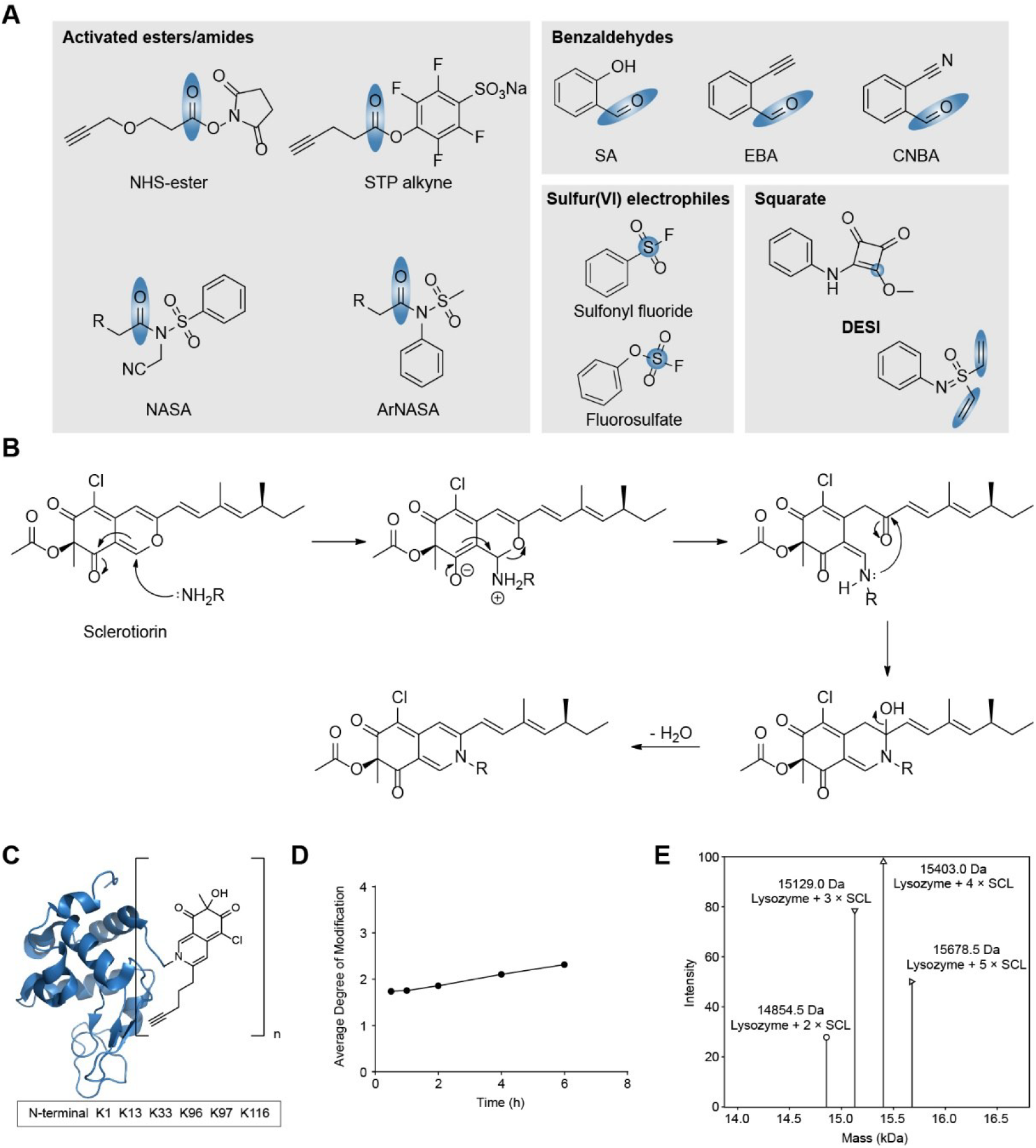
(A) Selected examples of previously reported chemistries for lysine targeting. Electrophilic sites susceptible to lysine attack are highlighted in blue. (B) Proposed reaction mechanism between sclerotiorin and a lysine residue. (C) Modification of lysozyme by SCL-alkyne.Lysozyme was selected due to the presence of six lysine residues and one *N*-termini. (D) Average degree of modification after incubation of lysozyme (0.5 mg/mL) with **SCL-alkyne** (7 eq.) in HEPES buffer (10 mM, pH 7.4) at 37 °C. The average degree of modification was calculated based on intact protein MS analysis. (E) Intact protein MS analysis of lysozyme after incubation with 7 equivalents of **SCL-alkyne** for 24 h. The intensity represents the deconvoluted mass signal obtained using UniDec.^[42]^

While many of these molecules are of synthetic origin, nature employs similar cyclization strategies for irreversible lysine trapping, including those found in the large class of azaphilones. Azaphilones such as **sclerotiorin** (**SCL**) were discovered decades ago as fungal metabolites with anti-inflammatory, cytotoxic, and antibacterial activities.^[35-36]^ They have attracted attention as lysine-modifying motifs for the introduction of protein modifications, for example as a suitable ligation strategy for antibody-drug conjugates.^[37]^ The hallmark of the azaphilone unit is a highly oxygenated bicyclic pyranoquinone ring that, upon nucleophilic attack by the lysine amino group, undergoes a cascade of ring-opening and ring-closing reactions to yield a vinylogous γ-pyridone (Figure 1B).^[37]^ Recently, it was shown that the core unit required to catalyze this trapping step can be reduced to a 3-acyl-4-pyranone scaffold, which is sufficient to capture lysine residues.^[38]^ Despite this lysine-tailored chemistry, studies deciphering the selectivity and extent of lysine liganding in whole proteomes, as well as the target scope of natural product-derived azaphilones, are still lacking.

Here, we synthesized a simplified sclerotiorin-derived probe (**SCL-alkyne**) that displays high selectivity for lysine residues, while achieving broad coverage of the *Escherichia coli* (*E. coli*) K12 proteome, capturing around 6,000 lysine sites, which is comparable to the benchmark probe **STP-alkyne**. Global quantitative profiling with **SCL-alkyne** enabled a systematic assessment of lysine reactivity and revealed that only a limited subset of engaged hyperreactive lysines corresponded to annotated functional sites. Among the enriched protein targets, several were associated with bacterial chemotaxis, which was confirmed by the reduced response of compound-treated bacteria towards a glucose-filled capillary. Beyond establishing **SCL-alkyne** as a broadly applicable tool for lysine-directed proteome profiling, we leveraged its structural resemblance to natural azaphilones to investigate the target scope of this compound class and identified iron-sulfur cluster assembly and antioxidant proteins as major targets.

## Results

### Synthesis of a sclerotiorin-derived probe and investigation of its *in vitro* lysine selectivity

Given the high structural diversity of azaphilone natural products, we selected **SCL** because of its reported bioactivities and well-established synthetic accessibility (Figure 1B).^[37]^ To investigate azaphilone-based proteome reactivity, we designed an alkyne-functionalized **SCL** probe by introducing a clickable handle into the aliphatic side chain (Scheme 1). This probe, termed **SCL-alkyne**, enables irreversible target enrichment followed by copper-catalyzed azide-alkyne cycloaddition (CuAAC) with biotin azide, carboxylate bead-based cleanup, streptavidin enrichment, trypsin digestion, and LC-MS/MS analysis for the identification of probe-bound proteins (Figure 2A).^[39-40]^ In addition, the use of isotopically labeled desthiobiotin azide (isoDTB) tags enables not only the quantitative assessment of lysine engagement, but also the ranking of modified lysine residues according to their relative reactivity (Figure 3A, S1).^[41]^

**Figure 2.**
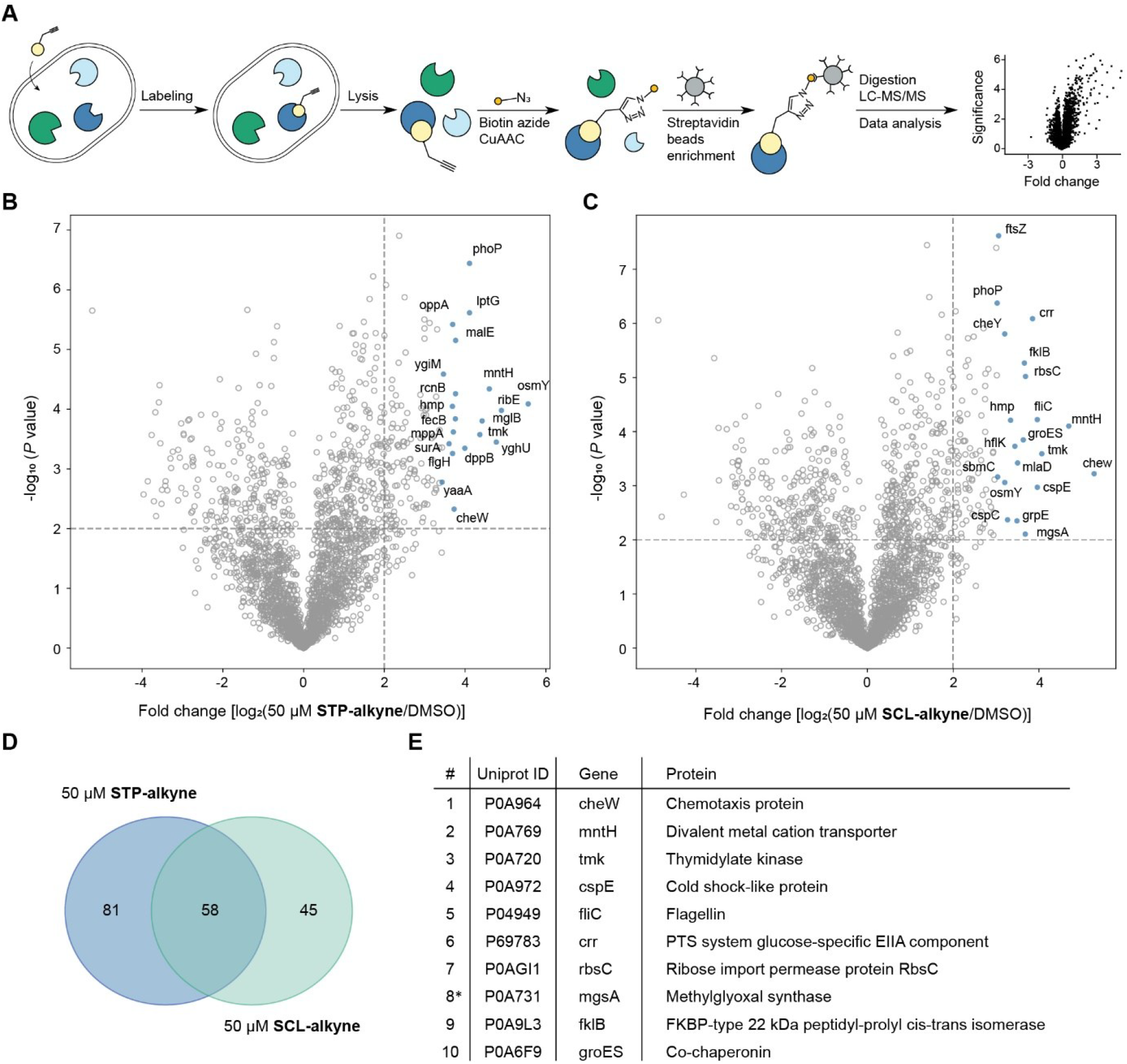
(A) Workflow for protein target identification using ABPP. Live *E. coli* K12 cells were incubated with **SCL-alkyne, STP-alkyne** or DMSO as a control. Probe-labeled proteins were subsequently enriched *via* biotin-based enrichment and digested prior to LC-MS/MS analysis. (B, C) Volcano plots showing proteins enriched after treatment with (B) **STP-alkyne** (50 μM, 1 h) or (C) **SCL-alkyne** (50 μM, 1 h). Thresholds were set at log_2_ fold change > 2.0 and p-value < 0.01 (two-tailed Student’s t-test, n = 4 replicates per group). The top 20 proteins most enriched by **STP-alkyne** or **SCL-alkyne** were annotated with their corresponding gene names. See Table S1 for the list of identified proteins. (D) Venn diagram of proteins enriched by **STP-alkyne** and **SCL-alkyne** at 50 μM, respectively. (E) The top 10 proteins most enriched by **SCL-alkyne**, including their ranking numbers, UniProt IDs, gene names, and protein names. *Asterisks indicate proteins uniquely enriched by SCL-alkyne. All others are overlapping proteins.

**Figure 3.**
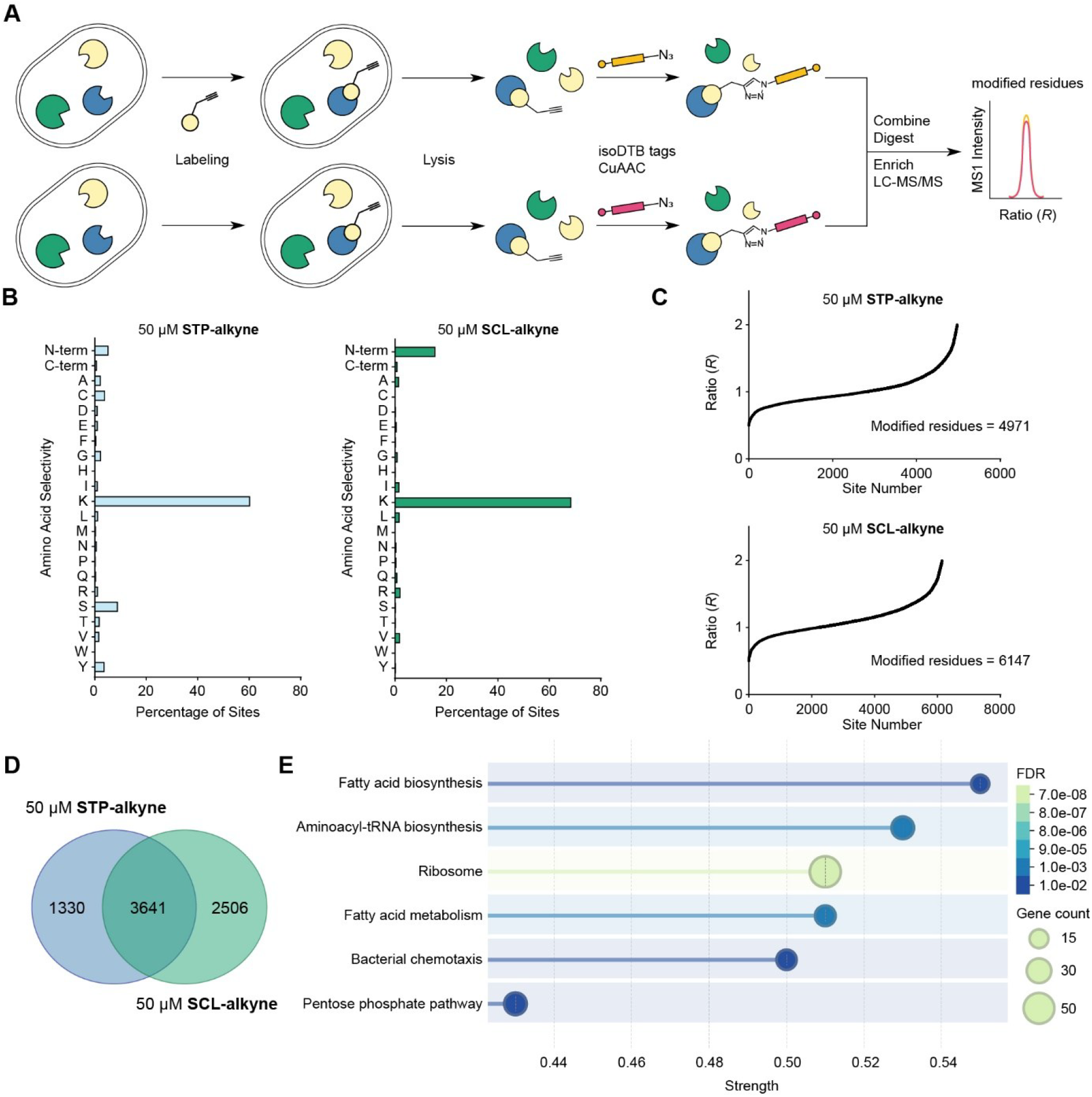
(A) Workflow for the identification of engaged amino acid residues using isoDTB tags. Following conjugation with isotopic tags, enrichment and digestion, labeled peptides were subjected to LC-MS/MS analysis and quantified according to their MS1 signal intensities. Amino acid sites targeted by the probe showed ratios (*R*) close to 1 between the heavy and light channels. (B) The bar graphs show the distribution of modified amino acid residues, including *C*-terminal (C-term) and *N*-terminal (*N*-term) modifications, in *E. coli* K12 cells treated with 50 μM **STP-alkyne** or **SCL-alkyne.** Residue assignments were based on differential modification analysis across four technical replicates. See Table S3 for the amino acid selectivity data. (C) *R* values of lysine residues labeled by **STP-alkyne** and **SCL-alkyne** at 50 μM in *E. coli* K12 cells were determined from four technical replicates. Only lysine sites with *R* values between 0.5 and 2 were considered modified residues. See Table S4 for the lysine-residue quantification data. (D) Venn diagram of engaged lysine sites identified by **STP-alkyne** and **SCL-alkyne** at 50 μM. (E) KEGG pathway analysis with 50 μM **SCL-alkyne** binding proteins from closed search in the above isoDTB workflow. This data was analyzed *via* String with Group Similarity ≥ 0.8 and medium confidence ≥ 0.4. The strength value reflects the magnitude of enrichment relative to the number of labeled and expected annotated proteins.^[43]^

As shown in Scheme 1, **SCL-alkyne** was synthesized from precursor **1** through benzoylation, oxime formation, bromination, and deprotection to furnish brominated aldehyde **6**. Through a Sonogashira reaction, the alkyne side chain was coupled to afford intermediate **7**, from which oxidative cyclization delivered compound **8**. However, compound **8** proved unstable and gradually hydrolyzed at room temperature. Therefore, chlorination of **7** yielded intermediate **9**, which following oxidative cyclization, furnished **SCL-alkyne**.

**Scheme 1.**
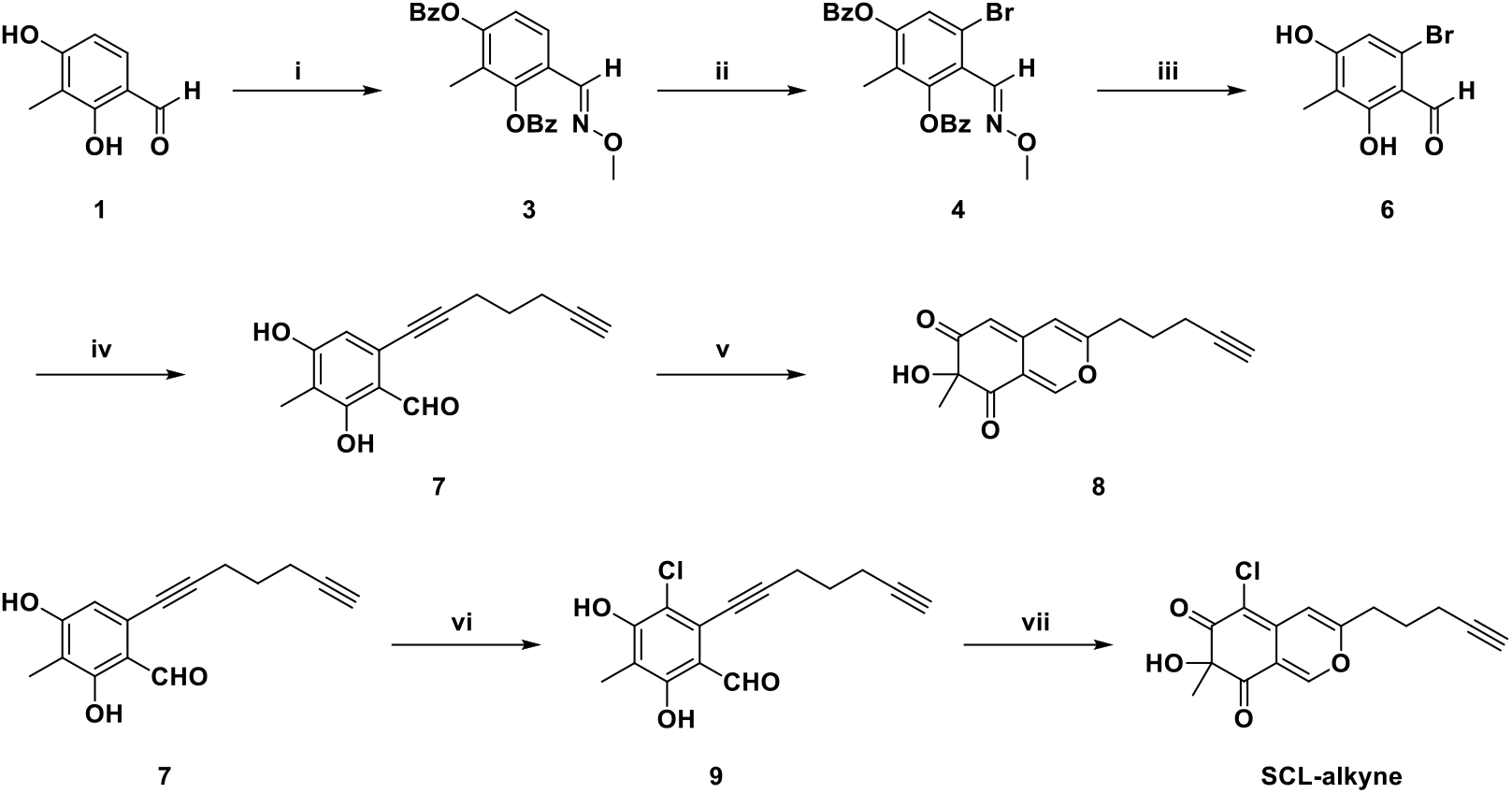
Synthesis of compound **8** and **SCL-alkyne**. Reagents and conditions: (i) (a) BzCl, Et_3_N, DMAP, DCM, rt, overnight, (b) CH_3_ONH_2_·HCl, Pyridine, DCM, rt, 1h, 82%; (ii) DBDMH, AcOH, CF_3_CO_2_Ag, Pd(OAc)_2_, DCE, 100 °C, 24h, 34%; (iii) (a) NaOH, MeOH, rt, overnight, (b) *p*-TsOH, CH_2_O, THF/H_2_O, 100 °C, overnight, 47%; (iv) 1,6-Heptadiyne, Pd(PPh_3_)_4_, CuI, Et_3_N, anhydrous THF, Ar, 80 °C, 20h, 22%; (v) (a) AgNO_3_, TFA, DCE, rt, 10 min, (b) IBX, TBAI, rt, 3h, 25%; (vi) SO_2_Cl_2_, DCM, rt; (vii) (a) AgNO_3_, TFA, DCE, rt, 10 min, (b) IBX, TBAI, rt, 3h, 34%.

**SCL-alkyne** was first tested against a panel of nucleophilic amino acids, including *N*(α)-acetyl lysine, *N*-acetyl cysteine, *N*(α)-acetyl arginine, *N*-acetyl tyrosine, and *N*-benzyloxycarbonyl serine, among which only incubation with lysine showed the corresponding mass shift of the anticipated γ-pyridone product (Figure S2). Based on this result, we next examined its reactivity toward lysozyme, a model protein containing six lysine residues and one *N*-termini, by incubation with seven equivalents of **SCL-alkyne** (Figure 1C). Time-dependent modification of lysine residues was monitored over 6 h, with the first labeling event occurring within 30 min (Figure 1D). Notably, up to five lysine residues were found to be modified at 24 h, suggesting differences in lysine reactivity depending on their location within the protein structure (Figure 1E).

### The azaphilone moiety selectively modifies lysine residues in live cells

With lysine selectivity confirmed on isolated amino acids, we commenced with whole-proteome labeling studies. Before performing target profiling of **SCL-alkyne** in live *E. coli* K12 cells, we first evaluated its antibacterial activity. **SCL-alkyne** showed no antibacterial activity against *E. coli* K12 at concentrations up to 1 mM. For analytic labeling, intact *E. coli* K12 cells were incubated with 50 μM **SCL-alkyne** or **STP-alkyne**, a well-established lysine-labeling reagent, followed by cell lysis and click reaction with rhodamine azide. Fluorescent SDS-PAGE analysis revealed labeling of multiple proteins with a slightly different pattern of the two probes and a higher intensity in case of **STP-alkyne** (Figure S3). We next compared the proteins labeled by **STP-alkyne** and **SCL-alkyne** following click chemistry with biotin azide, enrichment on streptavidin-coated beads, digestion and LC-MS/MS analysis (Figure 2A). At the proteome level, the labeling profiles of the two compounds were broadly comparable (Figure 2B, 2C). A considerable overlap was observed, with 58 proteins enriched by both probes, whereas 81 and 45 proteins were uniquely enriched by **STP-alkyne** and **SCL-alkyne**, respectively, demonstrating that **SCL-alkyne** exhibits proteome-labeling performance comparable to that of a well-established lysine-labeling reagent, while also covering a complementary set of targets (Figure 2D). Notably, 9 of the top 10 most enriched proteins identified by **SCL-alkyne** were found within the overlap area, suggesting that their labeling is primarily driven by highly reactive lysine sites, rather than by the unique structural features of **SCL-alkyne** (Figure 2E).

To further identify the corresponding lysine modification sites on the labeled proteins, we applied the above-described isoDTB workflow using **SCL-alkyne**, with **STP-alkyne** as a benchmark control (Figure 3A). In an Open Search, mass-shift analysis confirmed that both probes reacted cleanly in the proteome. Apart from unmodified peptides and common background modifications such as carbamidomethylation (57.0215 Da) and oxidation (15.9949 Da), the mass shift was dominated by the expected probe-derived modifications (Figure S4). These mass shifts corresponded to probe-modified peptides after conjugation with the light/heavy tags, namely 755.3166/761.3226 Da for **SCL-alkyne** and 561.3014/567.3108 Da for **STP-alkyne** (Table S2). To assess amino acid selectivity, an Offset Search was performed, revealing that 84% of all **SCL-alkyne**-modified sites at 50 μM were assigned to lysine residues and *N*-termini, with negligible labeling of other amino acids (Figure 3B, Table S3). This selectivity was even retained at concentrations up to 100 μM (Figure S5A). In contrast, **STP-alkyne** showed lower selectivity, with only 66% of modified sites corresponding to lysine residues and *N*-termini (Figure 3B, Table S3). Notably, **SCL-alkyne** enabled the localization of the modification to *N*-termini of 470 proteins at 100 μM, corresponding to a selectivity of approximately 15%, which is higher than that of **STP-alkyne** (5%), **ArSq-alkyne** (2%), and **EBA-alkyne** (5%) (Table S3).^[18]^ This finding may help address the current lack of chemistries with strong selectivity for *N*-termini.

Moreover, in a Closed Search, **SCL-alkyne** modified 6,147 lysines in 1,167 proteins at 50 μM and 7,690 lysines in 1,389 proteins at 100 μM throughout the proteome of intact *E. coli* K12 cells (Figure 3C, S5B, Table S4). By comparison, **STP-alkyne** modified 4,971 lysines in 1,003 proteins at 50 μM (Figure 3C). Although a substantially larger overlap was observed, **SCL-alkyne** still engaged 2,506 distinct lysine sites, indicating that it can serve as a complementary lysine labeling tool (Figure 3D). Furthermore, 766 of the **SCL-alkyne** labeled proteins were annotated as intracellular proteins, suggesting that the azaphilone scaffold is capable of penetrating the poorly permeable double membrane of *E. coli*, an important feature for bacterial applications (Figure S6). Given that the pK_a_ values of lysine ε-amino groups can vary depending on the local microenvironment, lysine-directed warheads that can cross the bacterial cell and react with lysine residues in intact live cells are particularly valuable. KEGG pathway analysis further revealed that many bound proteins are involved in fatty acid biosynthesis, aminoacyl-tRNA biosynthesis, ribosome function, fatty acid metabolism, and bacterial chemotaxis (Figure 3E).

The reactivity of lysine residues is known to vary within proteins depending on their local environment, for example through interactions with neighboring amino acids that directly influence pK_a_ values and nucleophilicity.^[44-45]^ Amino acid reactivity profiling has been established as a powerful strategy for identifying hyperreactive residues. Here, we similarly applied **SCL-alkyne** to profile lysine reactivity in *E. coli* at high (100 μM) and low (10 μM) concentrations (Figure S1). Highly reactive lysine sites are expected to become saturated already at the lower concentration and, therefore, display an isotopic ratio close to 1 (Figure 4A). In contrast, less reactive lysine sites showed progressively higher ratios depending on their intrinsic reactivity. To exclude false quantification events, only lysines detected in the control experiment with isotopic ratios between 0.5 and 2, were considered for further analysis (Table S5). Using this approach, we quantified 4,935 lysines and identified 79 highly reactive lysines (*R*_10:1_ < 3) in 72 different proteins. An additional 260 lysines displayed medium reactivity (3 < *R*_10:1_ < 5) in 217 proteins, whereas the remaining 4,596 lysines were classified as low-reactivity sites (*R*_10:1_ > 5) in 1,023 proteins (Figure 4B). Regarding the distribution of reactive lysine sites, approximately 85% of proteins with a high or medium reactive lysine contained more than one quantified lysine residue, whereas nearly 80% of these proteins harbored only a single high or medium reactive lysine site (Figure 4D).

**Figure 4.**
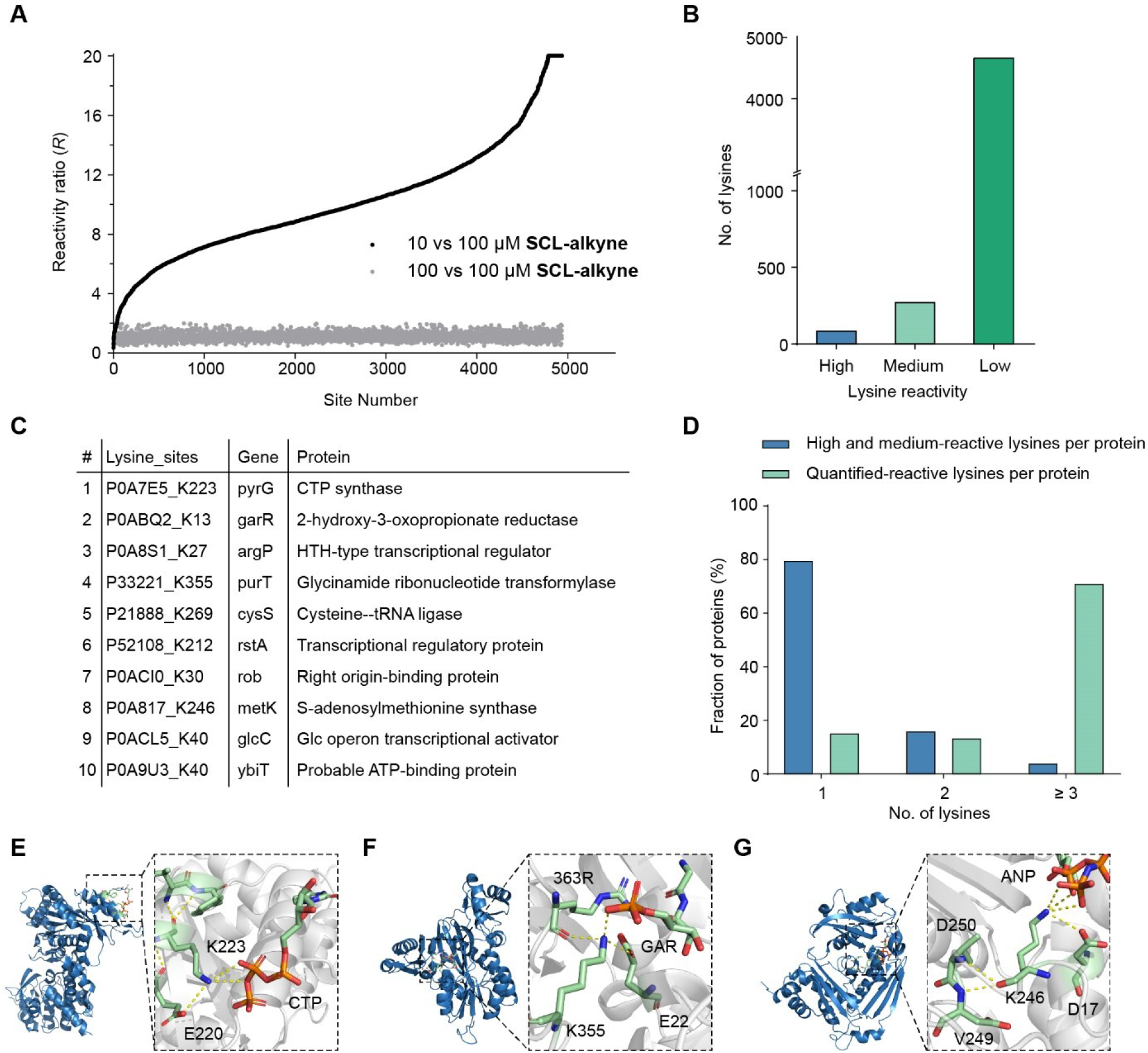
(A) Plot of the reactivity ratios (*R*_10:1_) obtained by comparing *E. coli* K12 cells treated with high (100 μM) vs. low concentration (10 μM) of **SCL-alkyne** (black). Ratios (*R*_1:1_) of an experiment with high concentration used for both samples (grey) are used as a control to ensure reliable quantification of all lysines. All data are based on four technical replicates. For visualization, *R*_10:1_ > 20 are shown as 20 in the plot. The corresponding original values are provided in Table S5. (B) Number of lysines in the different reactivity bins, classified as highly reactive (*R*_10:1_ < 3), medium-reactivity (3 < *R*_10:1_ < 5), and low-reactivity (*R*_10:1_ > 5) based on their isotopic ratios. (C) Top 10 most reactive lysine sites ranked by isotopic ratio that have a direct functional annotation, including UniProt ID, lysine position, gene name, and protein name. (D) Percentage of high and medium reactive lysines, as well as quantified lysine sites, per protein with at least one high or medium reactive lysine. (E, F, G) Crystal structures of PyrG (PDB: 2AD5), PurT (PDB: 1EZ1), and MetK (PDB: 1P7L), as well as magnified views of the interactions between their engaged lysine sites and neighboring amino acid residues. CTP: cytidine triphosphate; GAR: glycinamide ribonucleotide; ANP: adenylyl imidodiphosphate. Interactions between residues were analyzed using PyMOL.

We next examined the top 10 lysine sites exhibiting the lowest isotopic ratios (Table S5). Structural analysis based on either resolved X-ray crystal structures or AlphaFold3 predictions revealed that these top 10 engaged lysine residues are located on solvent-accessible surface area. We therefore hypothesize that the reduced steric constraints of these residues contribute to their elevated intrinsic reactivity. We further analyzed engaged lysine sites located within five amino acid residues of catalytic sites, substrate-binding sites, DNA-binding sites, or other annotated functional regions according to UniProt annotations (Table S5). In total, 436 engaged lysine sites were identified in proximity to these functional regions. Notably, only 6.7% of these sites showed high or medium reactivity. Within this proximal subset, 130 sites directly corresponded to annotated functional sites, of which 9.2% showed high or medium reactivity (Table S5).

Because most functionally annotated engaged lysines were not classified as highly reactive sites in the reactivity profiling experiment, we did not restrict the subsequent structural analysis to a small subset. Instead, to identify common structural characteristics among functionally annotated engaged lysines, we focused on the 130 residues that directly corresponded to annotated sites, regardless of their reactivity class (Figure 4C, Table S5). These included several lysines in established drug targets, such as GyrB (K103 in the DNA gyrase subunit B, UniProt ID: P0AES6), DdlA (K185 in D-alanine--D-alanine ligase A, UniProt ID: P0A6J8), AccC (K202 in Acetyl-CoA carboxylase biotin carboxylase subunit, UniProt ID: P24182) and NrdA (K91 in Ribonucleoside-diphosphate reductase subunit alpha, UniProt ID: P00452). Interestingly, most of these sites formed hydrogen bonds or potential electrostatic interactions with neighboring aspartate or glutamate residues. For example, K223 in cytidine triphosphate (CTP) synthetase forms a hydrogen bond with the triphosphate moiety while simultaneously interacting with E220 through an additional hydrogen bond (Figure 4E). Similarly, the binding sites K355 in PurT transformylase and K246 in S-adenosylmethionine synthetase (MetK) also interact with neighboring aspartate or glutamate residues (Figure 4F, 4G). The interactions between lysine residues and neighboring acidic residues may help fine-tune the lysine protonation state, while also limiting excessive intrinsic reactivity. Thus, the identification of ligandable lysine sites may require a more comprehensive evaluation integrating intrinsic reactivity and structural environment.

### Proteome profiling identifies proteins involved in bacterial chemotaxis

Natural products bearing an azaphilone moiety display remarkable structural diversity and broad biological activity. To gain insight into their native biological functions and to directly interrogate their cellular targets, we applied **SCL-alkyne** to intact *E. coli* K12 cells and profiled labeled proteins by protein-level ABPP (Figure 2A). Labeling was performed at probe concentrations of 10, 50, and 100 μM (Figure 2C, S7, Table S1). Because only a limited number of proteins were labeled at 10 μM, our analysis primarily focused on the overlap between the 50 and 100 μM conditions (Figure 5A). In total, 100 proteins were enriched under both conditions, many of which are associated with bacterial chemotaxis and ATP-binding cassette (ABC) transporters. Consistently, these two functional terms were also significantly enriched in the KEGG pathway analysis (Figure S8). Notably, several key chemotaxis proteins, including CheW and CheY, and MglB, MalE, two periplasmic solute-binding proteins that bridge nutrient transport and chemotaxis, emerged as strongly enriched hits. The corresponding binding sites had already been identified in the isoDTB workflow. The most reactive sites within these proteins were K56 in CheW, K26 in CheY, K332 in MglB, and K303 in MalE (Table S5). Among them, CheY K26 and MalE K303 were classified as highly reactive lysine sites (Table S5). Although the corresponding binding sites exhibited different levels of reactivity, their enrichment intensities were roughly comparable (Figure 5B).

**Figure 5.**
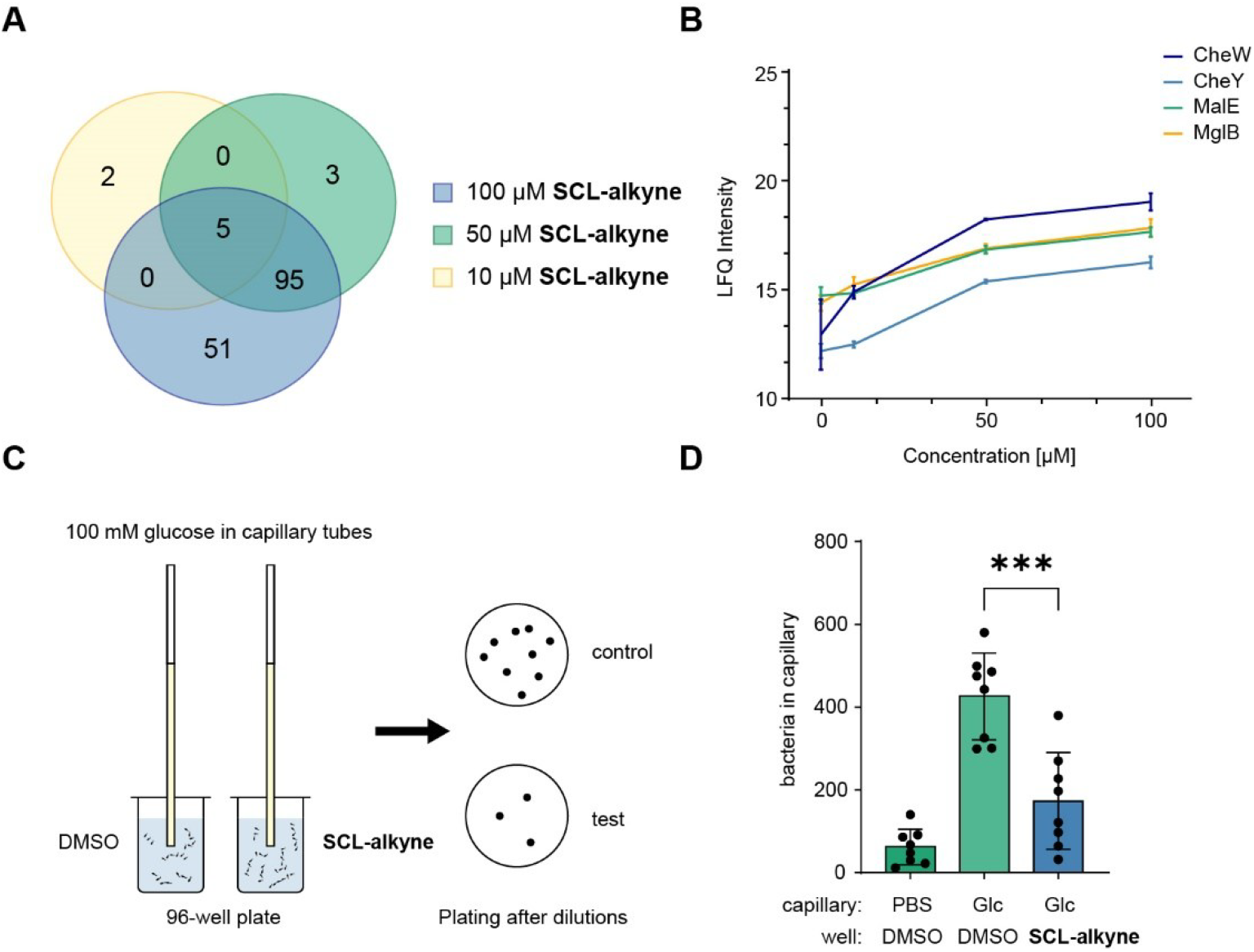
(A) Venn diagram showing the overlap of proteins significantly enriched at 100, 50, and 10 μM probe concentrations. (B) Dose-dependent Label-Free Quantification (LFQ) intensities of chemotaxis-associated proteins (two-tailed Student’s t-test, n = 4 replicates per group). (C) Workflow of the capillary chemotaxis assay. Bacteria swim from a reservoir in a 96-well plate into a glass capillary filled with an attractant, 100 mM glucose. After 60 min, the number of bacteria in the capillary is determined by plating the cells on agar plates. (D) Results of the capillary chemotaxis assay. The number of bacteria was determined by counting colonies on agar plates. The strain used here was *E. coli* K12. Glc, glucose. Data are shown as mean ± SD from eight biological replicates (n = 8). Statistical significance was determined using one-way ANOVA. (***p ≤ 0.001).

To examine whether engagement of chemotaxis-related proteins affects *E. coli* motility, we first performed a microscopy-based tracking assay. This analysis revealed no measurable differences in either swimming distance or swimming velocity upon treatment with **SCL-alkyne**, indicating that the compound does not directly impair bacterial motility (Figure S9). We therefore considered the alternative possibility that **SCL-alkyne** perturbs chemotactic behavior rather than the motility itself. To test this, we conducted a capillary chemotaxis assay (Figure 5C). In this assay, the *cheY* or *cheW* knockout strains no longer responded to the sugar stimulus, confirming that altered bacterial behavior in this setup reflects disruption of the chemotaxis system (Figure S10).^[46]^ Strikingly, treatment with 100 μM **SCL-alkyne** produced a similar phenotype, supporting the hypothesis that the compound interferes with bacterial chemotaxis (Figure 5D). The ability of **SCL-alkyne** to penetrate Gram-negative bacteria, covalently engage intracellular targets, and attenuate the bacterial response to environmental cues represents an unprecedented finding. These results highlight a potentially sophisticated evolutionary strategy by which azaphilone natural products modulate bacterial behavior through selective lysine liganding. They also demonstrate the utility of ABPP, as the involvement of the chemotaxis system would have been difficult to elucidate without comprehensive analysis of the enriched proteins.

### Competitive profiling of natural azaphilones reveals a variety of targeted cellular pathways

Intrigued by the wealth of **SCL-alkyne** targets in *E. coli*, we next extended our analysis to a panel of mitorubrin natural products (**mitorubrin, mitorubrinol** and **mitorubrinol acetate**) (Figure 6A). These compounds all contain an azaphilone scaffold bearing phenolic moieties. To date, little is known about their cellular targets or mechanisms-of-action, and target identification studies may, therefore, provide important insights into both their bioactivities and their underlying modes-of-action. However, the structural complexity of these molecules poses a substantial challenge for the synthesis of individual chemical probes. Thus, based on the broad lysine reactivity of **SCL-alkyne**, resembling a minimal azaphilone scaffold, we rationalized that competitive profiling on the protein level with structurally more complex analogs could reveal their target preferences, while also validating the probe as a general tool for assessing lysine reactivity.

**Figure 6.**
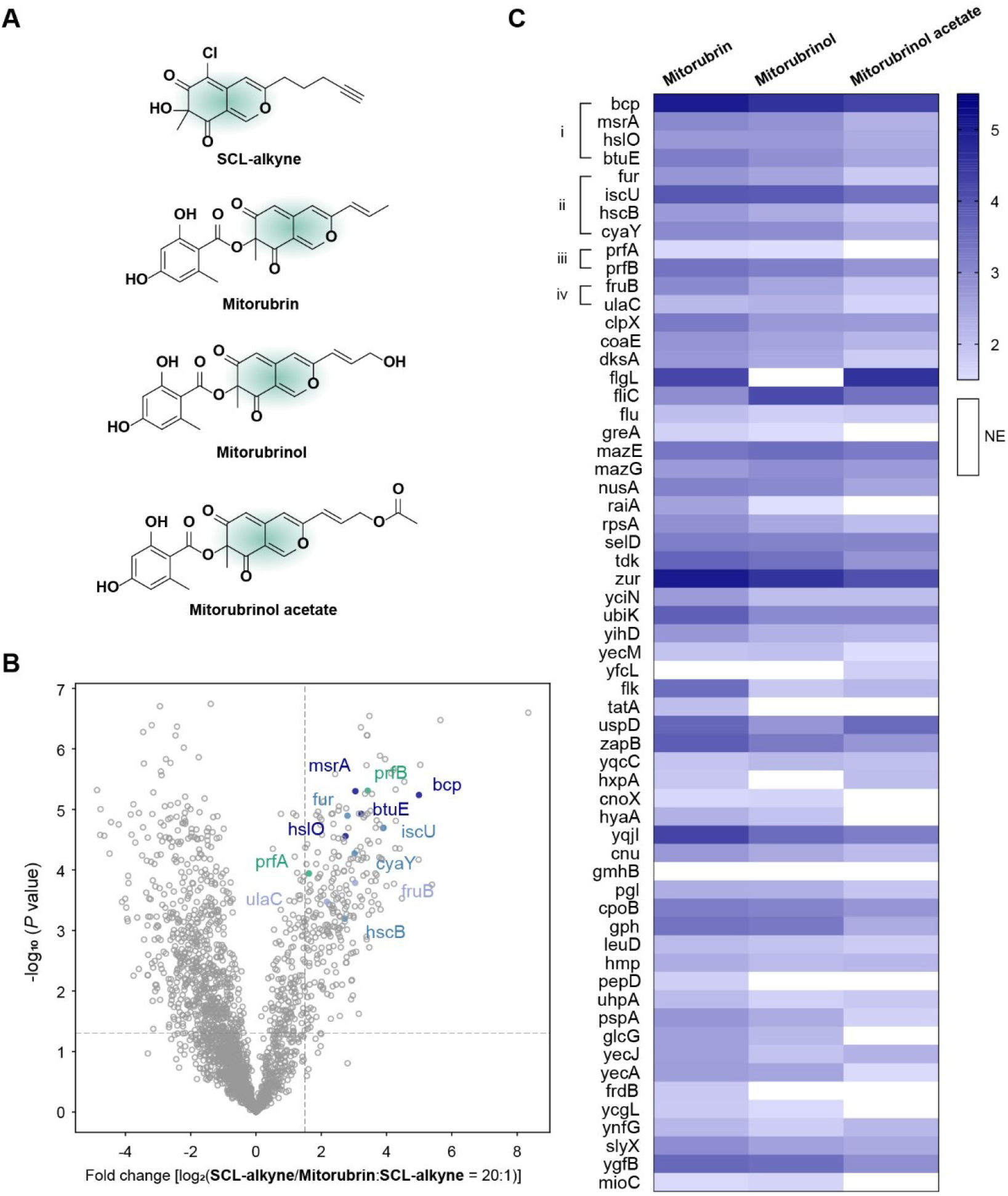
(A) Structures of **SCL-alkyne, mitorubrin, mitorubrinol** and **mitorubrinol acetate.**(B) Volcano plot of competitive ABPP in intact *E. coli* K12 cells. The plot shows preparative labeling results of cells treated with **SCL-alkyne** alone (50 μM, 1 h) compared to **SCL-alkyne** labeling in the presence of 1 mM **mitorubrin**. Thresholds were set at log_2_ fold change > 1.5 and p-value < 0.05 (two-tailed Student’s t-test, n = 4 replicates per group). Proteins belonging to the clusters identified in the heat map were annotated with their gene names and color-coded according to biological process. Dark blue: oxidative stress; light blue: iron-sulfur cluster assembly; green: protein-containing complex disassembly; light purple: phosphorylation. See Table S6 for the list of identified proteins. (C) Heat map showing competition ratios of proteins enriched by **SCL-alkyne** and competed by at least one of the three mitorubrin natural products. Competition ratios were calculated as **SCL-alkyne** labeling relative to **SCL-alkyne** labeling after preincubation with each mitorubrin natural product. Four major clusters were identified: (i) response to oxidative stress, (ii) iron-sulfur cluster assembly, (iii) protein-containing complex disassembly, and (iv) phosphorylation. NE, not enriched (fold change < 1.5).

To this end, competitive profiling was performed by preincubating intact *E. coli* K12 cells with each natural product at 1 mM prior to addition of the probe (mitorubrins exhibit no antibiotic activity under these conditions). Following cell lysis and the standard enrichment workflow, competed targets were identified by LC-MS/MS analysis (Figure 6B, S11). In total, 60 protein targets were found to be outcompeted by one or more of the natural products (Figure 6C). Based on these findings, we performed enrichment analysis of biological process, which revealed that many of the competed proteins are associated with iron-sulfur cluster assembly and antioxidant activity (Figure 6C). As all mitorubrins exhibit phenolic groups as putative iron chelators, we observed immediate adduct formation and visible precipitation upon mixing **mitorubrin** with iron (Figure S12). Additional studies will be required to clarify how the natural products influence these cellular pathways along with the biological relevance of their apparent interaction with iron.

## Conclusion

In summary, using the probe **SCL-alkyne**, we demonstrate that the azaphilone scaffold shows high selectivity toward lysine residues and *N*-termini in the proteome, while enabling broad proteome coverage in intact *E. coli* cells. This probe allowed systematic profiling of lysine reactivity on a proteome-wide scale, revealing that only a relatively small subset of lysines exhibits high or moderate intrinsic reactivity towards azaphilones. Notably, some of these sites are located in functionally important regions, including catalytic and cofactor-binding sites, thereby supporting the notion that lysine-directed covalent liganding represents a tractable strategy for ligand design against selected proteins, particularly those containing reactive lysines within catalytic pockets.

Beyond its value as a global lysine-profiling reagent, **SCL-alkyne** also provided direct insight into the biological role of azaphilone-based structures. Proteome-wide enrichment analysis identified chemotaxis-related proteins as prominent targets, and functional assays showed that probe treatment perturbs bacterial chemotactic behavior without measurably affecting motility itself. Together, these findings suggest that azaphilone electrophiles can modulate bacterial responses to environmental stimuli by selectively targeting intracellular lysines on chemotaxis-related proteins, while also highlighting the ability of this scaffold to penetrate Gram-negative bacteria.

Importantly, the broad proteome coverage of **SCL-alkyne** could be further leveraged in a competitive chemoproteomic strategy for structurally more complex azaphilone natural products. Without the need for individual probe synthesis, this approach enabled the identification of their cellular targets and revealed overlapping target pathways, including proteins associated with iron-sulfur cluster assembly and oxidative stress-related processes. Taken together, our findings position the azaphilone as a promising scaffold for the development of lysine-directed covalent probes and inhibitors, while also providing a broadly applicable strategy for elucidating the targets and biological functions of azaphilone natural products.

### Associated content

The mass spectrometry proteomics data have been deposited to the ProteomeXchange Consortium *via* the PRIDE partner repository with the dataset identifier PXD079507. Additional experimental details, including procedures, figures, tables, and spectra, are provided in the Supporting Information.

## Author information

Corresponding Authors

Stephan A. Sieber – Center for Functional Protein Assemblies (CPA), Department of Bioscience, TUM School of Natural Sciences, Technical University of Munich (TUM), Ernst-Otto-Fischer-Straße 8, 85748 Garching, Germany. https://orcid.org/0000-0002-9400-906X;

### Authors

Wei Ding – Center for Functional Protein Assemblies (CPA), Department of Bioscience, TUM School of Natural Sciences, Technical University of Munich (TUM), Ernst-Otto-Fischer-Straße 8, 85748 Garching, Germany.

Sophie Brameyer – Faculty of Biology, Microbiology, Ludwig Maximilian University of Munich, 82152 Martinsried, Germany.

Susanne H. Kirsch – Department of Microbial Drugs, Helmholtz Centre for Infection Research (HZI), Helmholtz Institute for Pharmaceutical Research Saarland (HIPS), University Campus E 8.1, 66123 Saarbrücken, Germany and German Centre for Infection Research (DZIF), DZIF Partner Site Hannover-Brunswick, Germany, Inhoffenstrasse 7, 38124, Brunswick, Germany.

Birthe Sandargo – Department of Microbial Drugs, Helmholtz Centre for Infection Research (HZI) and German Centre for Infection Research (DZIF), DZIF Partner Site Hannover-Brunswick, Germany, Inhoffenstrasse 7, 38124, Brunswick, Germany.

Frank Surup – Department of Microbial Drugs, Helmholtz Centre for Infection Research (HZI) and German Centre for Infection Research (DZIF), DZIF Partner Site Hannover-Brunswick, Germany, Inhoffenstrasse 7, 38124, Brunswick, Germany. Current address: Department of Molecular Structural Biology, Helmholtz Centre for Infection Research, 38124 Braunschweig, Germany.

Stephan M. Hacker – Department of Molecular Physiology, Leiden Institute of Chemistry, Leiden University, Einsteinweg 55, 2333 CC, Leiden, The Netherlands.

Kirsten Jung – Faculty of Biology, Microbiology, Ludwig Maximilian University of Munich, 82152 Martinsried, Germany.

### Notes

The authors declare the following competing financial interest(s): S.A.S. is cofounder of smartbax limited.

## Acknowledgements

This work was supported by the European Union, ERC, breakingBAC (Grant No. 101096911), program of China Scholarship Council (Grant No. 202206210146) and the Deutsche Forschungsgemeinschaft (DFG, German Research Foundation. Grant No. EXC3092/1- 533751719). S.M.H. acknowledges the Dutch Research Council (NWO) for funding through a VIDI grant (VI.Vidi.213.057). Susanne H. Kirsch, Birthe Sandargo, and Frank Surup were partially funded by the German Centre for Infection Research (DZIF) projects TTU 09.721 and TTU 09.826. We sincerely thank Prof. Dr. Marc Stadler for generously providing mitorubrins and other natural products from the DZIF Natural Compound Library, which were originally isolated from *H. fragiforme* specimens.^[47]^ We also acknowledge Kora Holschbach for her input on synthetic route exploration. We thank Dr. Leeroy Baron (TUM) for proofreading the manuscript. We are grateful to Dr. Nina Bach, Mona Wolff, and Katja Bäuml (CPA) for their technical support.

