## Supporting Information for "Natural product-derived azaphilones are selective scaffolds for covalent lysine liganding in live cells"

Note: The mass spectrometry proteomics data have been deposited to the ProteomeXchange Consortium (http://proteomecentral.proteomexchange.org) *via* the PRIDE partner repository with the dataset identifier PXD079507.[1],[2]

### Table of Contents

**1. Chemistry Methods** ... 4

1.1. General Information. ... 4

1.2. Synthetic Procedures[3],[4] ... 5

**2. Biological Methods** ... 8

2.1. Bacterial Culture Conditions ... 8

2.2. *In vitro* Modification of Amino Acids and Proteins by **SCL-alkyne**[4] ... 8

2.3. Gel-based Fluorescent *In situ* Labeling ... 8

2.4. Preparative Labeling with **SCL-alkyne** and **STP-alkyne**[7],[8] ... 9

*In situ* labeling and lysis ... 9

Click and enrichment ... 9

Digestion and desalting ... 10

2.5. Competitive Labeling with **Mitorubrin**, **Mitorubrinol** and **Mitorubrinol Acetate** ... 10

2.6. LC-MS/MS Measurements on TimsTOF Pro and Proteomic Data Analysis ... 10

Measurement parameters on timsTOF Pro ... 10

Proteomic data analysis ... 11

Volcano plot visualization ... 11

2.7. isoDTB Experiments for Amino Acid Selectivity, Lysine-Residue Quantification[11] ... 12

*In situ* labeling and lysis ... 12

Click with isoDTB tags and digestion ... 12

Enrichment and desalting ... 13

2.8. Lysine-Residue Reactivity Profiling ... 13

2.9. LC-MS/MS Analysis of isoDTB Samples on TimsTOF Pro ... 14

2.10. isoDTB Samples Data Analysis[12] ... 14

General setup of analysis software ... 14

Analysis of the mass of modifications with FragPipe (open search) ... 14

Analysis of amino acid selectivity with FragPipe (offset search) ... 15

Lysine-residue quantification with FragPipe (closed search) ... 16

2.11. Analysis of Functional Sites ... 18

2.12. Microscopic Motility Tracking Assay[16] ... 18

2.13. Chemotaxis Capillary Assay[17] ... 19

#### 1. Chemistry Methods

1.1. General Information.

Reagents were purchased from BLDpharm or Sigma-Aldrich and used without further purification. Anhydrous solvents, deuterated solvents, and HPLC-grade solvents were obtained from Sigma-Aldrich, VWR Chemicals, or Fisher Chemicals. Reactions sensitive to air or moisture were performed under argon using standard Schlenk techniques and flame-dried glassware or sealed tube. TLC analysis was carried out on aluminium-backed silica gel 60 F254 plates (Merck KGaA). Compounds were visualized by UV light at 254 nm. Flash column chromatography was performed on silica gel 60 (40-63 µm, 230-400 mesh ASTM, Merck KGaA).

Chemical structures and reaction schemes were prepared using ChemDraw Professional 20.0.0.41. High-resolution ESI mass spectra were acquired on an LTQ-FT Ultra or Q Exactive Plus mass spectrometer (Thermo Fisher Scientific) coupled to a Dionex Ultimate 3000 HPLC system. Data were analyzed using Xcalibur 2.2 software (Thermo Fisher Scientific). NMR spectra were acquired on Bruker Avance I or Avance III HD spectrometers and processed with TopSpin software (version 2.1/3, Bruker). Chemical shifts (δ) are given in ppm and referenced to residual solvent peaks: $CDCl_3$ at 7.26 ppm and DMSO-$d_6$ at 2.50 ppm for $^1H$ NMR; $CDCl_3$ at 77.16 ppm and DMSO-$d_6$ at 39.52 ppm for $^{13}C$ NMR. Multiplicities are indicated as s, singlet; d, doublet; t, triplet; q, quartet; quint, quintet; td, triplet of doublets; m, multiplet; and br, broad. Coupling constants (*J*) are reported in Hz. Spectra were analyzed using MestReNova software version 14.0.0-23239.

1.2. Synthetic Procedures[3],[4]

**(*E*)-4-((methoxyimino)methyl)-2-methyl-1,3-phenylene dibenzoate (3)**

To a solution of 2,4-dihydroxy-3-methylbenzaldehyde (2.00 g, 13.15 mmol) in 100 mL $CH_2Cl_2$, benzoyl chloride (3.06 mL, 3.70 g, 26.30 mmol), 4-dimethylaminopyridine (DMAP, 321 mg, 2.63 mmol), and triethylamine ($Et_3N$, 3.67 mL, 2.66 g, 26.30 mmol) were added at room temperature. The reaction mixture was stirred overnight. The reaction was then diluted with water and extracted with $CH_2Cl_2$ (3 × 50 mL). The combined organic phases were washed with brine, dried over anhydrous $Na_2SO_4$, and filtered. The solvent was removed under reduced pressure, and the crude product was used in the next step without further purification.

The crude product from the previous step (4.65 g, 12.90 mmol) was dissolved in 50 mL $CH_2Cl_2$. Methoxyamine hydrochloride (1.19 g, 14.19 mmol) and pyridine (4.16 mL, 4.08 g, 51.60 mmol) were added at room temperature. The reaction mixture was stirred for 1 h, diluted with water, and extracted with $CH_2Cl_2$ (3 × 20 mL). The combined organic phases were washed with brine, dried over anhydrous $Na_2SO_4$, and filtered. The solvent was removed under reduced pressure, and the crude product was purified by flash column chromatography (3% EtOAc in Hexane) to afford the product (4.20 g, 10.79 mmol, 82.05% over two steps) as an off-white solid. $^1$H NMR (500 MHz, Chloroform-*d*) δ 8.25 – 8.21 (m, 4H), 8.08 (s, 1H), 7.73 (d, *J* = 8.5 Hz, 1H), 7.69 – 7.65 (m, 2H), 7.56 – 7.52 (m, 4H), 7.18 (d, *J* = 8.5 Hz, 1H), 3.81 (s, 3H), 2.10 (s, 3H).

**(*E*)-5-bromo-4-((methoxyimino)methyl)-2-methyl-1,3-phenylene dibenzoate (4)**

To a solution of compound **3** (4.20 g, 10.79 mmol) in 100 mL 1,2-dichloroethane, 1,3-dibromo-5,5-dimethylhydantoin (DBDMH, 3.09 g, 10.79 mmol), $Pd(OAc)_2$ (243 mg, 1.08 mmol), silver trifluoroacetate (239 mg, 1.08 mmol), and acetic acid (0.62 mL, 648 mg, 10.79 mmol) were added at room temperature. The reaction mixture was stirred and refluxed at 100 °C for 24 h, filtered through Celite, and concentrated under reduced pressure. The crude product was purified by flash column chromatography (3% EtOAc in Hexane) to afford the product (1.70 g, 3.63 mmol, 33.64%) as an off-white solid. $^1$H NMR (500 MHz, Chloroform-*d*) δ 8.30 (s, 1H), 8.24 – 8.20 (m, 4H), 7.69 – 7.63 (m, 2H), 7.56 – 7.51 (m, 4H), 7.48 (s, 1H), 3.42 (s, 3H), 2.10 (s, 3H).

**6-bromo-2,4-dihydroxy-3-methylbenzaldehyde (6)**

To a solution of compound **4** (1.70 g, 3.63 mmol) in 50 mL MeOH, NaOH (2.90 g, 72.6 mmol) was added by portions at room temperature. The reaction mixture was stirred for 20 h. The reaction was then quenched with water in an ice bath, and the mixture was extracted with EtOAc (3 × 50 mL). The combined organic phases were washed with brine, dried over anhydrous $Na_2SO_4$, and filtered. The solvent was removed under reduced pressure, and the crude product was used in the next step without further purification.

The obtained crude compound (1.21 g, 4.65 mmol) was dissolved in THF/$H_2O$ (10:1, 50 mL). *p*-toluenesulfonic acid monohydrate (1.77 g, 9.30 mmol) and formaldehyde solution, 37 wt. % in water (3.47 mL, 46.50 mmol) were added, and the mixture was refluxed at 100 °C overnight. After cooling, the mixture was diluted with water and extracted with EtOAc (3 × 50 mL). The combined organic phases were washed with brine, dried over anhydrous $Na_2SO_4$, and filtered. The solvent was removed under reduced pressure, and the crude product was purified by flash column chromatography (5% EtOAc in Hexane) to furnish the desired compound (396 mg, 1.71 mmol, 47.11% over two steps) as an off-white solid. $^1$H NMR (500 MHz, Chloroform-*d*) δ 12.68 (s, 1H), 10.11 (s, 1H), 6.68 (s, 1H), 5.65 (s, 1H), 2.08 (s, 3H).

**6-(hepta-1,6-diyn-1-yl)-2,4-dihydroxy-3-methylbenzaldehyde (7)**

A suspension of compound **6** (150 mg, 0.65 mmol), 1,6-Heptadiyne (179 µL, 144 mg, 1.56 mmol), $Pd(PPh_3)_4$ (38 mg, 32.50 µmol), CuI (6 mg, 32.5 µmol), and triethylamine (272 µL, 197 mg, 1.95 mmol) in 5 mL of anhydrous THF was stirred at 80 °C in a sealed tube under argon for 20 h. The mixture was cooled to room temperature, and saturated $NH_4Cl$ solution was added. The mixture was extracted with EtOAc (3 × 10 mL), and the combined organic layers were washed with brine, dried over anhydrous $Na_2SO_4$, and filtered. The solvent was removed under reduced pressure, and the crude product was purified by flash column chromatography (5% EtOAc in Hexane) to afford the product (33 mg, 0.14 mmol, 21.54%) as a pale-yellow oil. $^1$H NMR (500 MHz, Chloroform-*d*) δ 12.33 (s, 1H), 10.20 (s, 1H), 6.49 (s, 1H), 2.60 (t, *J* = 7.0 Hz, 2H), 2.37 (td, *J* = 7.0, 2.5 Hz, 2H), 2.11 (s, 3H), 2.00 (t, *J* = 2.5 Hz, 1H), 1.84 (quint, *J* = 7.0 Hz, 2H).

**7-hydroxy-7-methyl-3-(pent-4-yn-1-yl)-6*H*-isochromene-6,8(7*H*)-dione (8)**

Compound **7** (33 mg, 0.14 mmol) was dissolved in 1,2-dichloroethane/TFA (10:1, 5 mL), and $AgNO_3$ (3 mg, 14 µmol) was added at room temperature. After stirring for 10 min, 2-iodoxybenzoic acid (IBX, 47 mg, 168 µmol) and tetra-*N*-butylammonium iodide (TBAI, 3 mg, 8.4 µmol) were added. The reaction mixture was stirred at room temperature for 3 h and then quenched with saturated $Na_2S_2O_3$ solution. The mixture was extracted with EtOAc (3 × 10 mL). The combined organic phases were washed with brine, dried over anhydrous $Na_2SO_4$, and filtered. The solvent was removed under reduced pressure, and the crude product was purified by flash column chromatography (1% to 3% MeOH in $CH_2Cl_2$) to afford the desired product (9 mg, 34.8 µmol, 24.86%) as a yellow oil. $^1$H NMR (500 MHz, Chloroform-*d*) δ 7.88 (d, *J* = 1.0 Hz, 1H), 6.16 (s, 1H), 5.53 (d, *J* = 1.5 Hz, 1H), 2.58 (t, *J* = 7.5 Hz, 2H), 2.30 (td, *J* = 6.5, 2.5 Hz, 2H), 2.02 (t, *J* = 2.5 Hz, 1H), 1.89 – 1.83 (m, 2H), 1.55 (s, 3H). $^{13}$C NMR (100 MHz, Chloroform-*d*) δ 196.48, 195.79, 161.86, 153.00, 143.94, 116.03, 109.03, 105.52, 83.67, 82.63, 70.01, 32.10, 28.64, 25.25, 17.85. HR-MS (ESI): m/z $[M+H]^+$ calcd for $C_{15}H_{15}O_4$: 259.0965; found: 259.0957.

**5-chloro-7-hydroxy-7-methyl-3-(pent-4-yn-1-yl)-6*H*-isochromene-6,8(7*H*)-dione (SCL-alkyne)**

Compound **7** (30 mg, 0.12 mmol) was dissolved in anhydrous $CH_2Cl_2$ (3 mL) and cooled to 0 °C. $SO_2Cl_2$ (12 µL, 20 mg, 0.15 mmol) was added, and the reaction mixture was warmed up to room temperature and stirred for 20 min. The reaction was carefully quenched with ice water and extracted with $CH_2Cl_2$ (3 × 10 mL). The combined organic phases were washed with brine, dried over anhydrous $Na_2SO_4$, and filtered. The solvent was removed under reduced pressure, and the crude product was used in the next step without further purification. In a manner similar to that used for compound **8**, **SCL-alkyne** was prepared from crude compound **9** and obtained as a pale-yellow oil (12 mg, 41 µmol, 34.17% over two steps). $^1$H NMR (500 MHz, Chloroform-*d*) δ 7.92 (s, 1H), 6.62 (s, 1H), 2.68 (t, *J* = 7.5 Hz, 2H), 2.33 (td, *J* = 6.5, 2.5 Hz, 2H), 2.05 (t, *J* = 2.5 Hz, 1H), 1.95 – 1.88 (m, 2H), 1.58 (s, 3H). $^{13}$C NMR (125 MHz, Chloroform-*d*) δ 194.07, 190.17, 163.94, 152.21, 139.62, 115.65, 109.05, 106.25, 84.19, 82.50, 70.18, 32.51, 28.73, 25.33, 17.94. HR-MS (ESI): m/z $[M+H]^+$ calcd for $C_{15}H_{14}ClO_4$: 293.0575; found: 293.0575.

#### 2. Biological Methods

##### 2.1. Bacterial Culture Conditions

Overnight cultures were prepared by inoculating 5 mL of medium with 5 µL of bacterial glycerol stock, followed by incubation at 37 °C and 200 rpm overnight. Unless stated otherwise, *E. coli* K12 was cultivated in Lysogeny Broth (LB; 10 g/L casein peptone, 5 g/L NaCl, and 5 g/L yeast extract). *E. coli* BW2511 Δ*cheY* and Δ*cheW* mutant strains were grown in LB medium supplemented with kanamycin at 25 µg/mL.[5]

##### 2.2. *In vitro* Modification of Amino Acids and Proteins by **SCL-alkyne**[4]

Amino acids were reacted with **SCL-alkyne** by incubating the amino acid (1.0 mM) with **SCL-alkyne** (2.0 mM) in phosphate buffer (100 mM, pH 8.0) at 37 °C for 1 hour. For peptide modification, lysozyme (0.5 mg/mL) was incubated with **SCL-alkyne** (7 equiv) in HEPES buffer (10 mM, pH 7.4) at 37 °C for the indicated time. All samples were measured on a Q Exactive Plus mass spectrometer (Thermo Fisher Scientific) coupled to a Dionex Ultimate 3000 HPLC system. Data acquisition was performed in positive ion mode with full MS scans at a resolution of 140,000 across an m/z range of 500-2000. Raw data were processed using Xcalibur 2.2 software (Thermo Fisher Scientific), and intact protein spectra were deconvoluted with UniDec 2.6.7.[6]

##### 2.3. Gel-based Fluorescent *In situ* Labeling

An overnight culture of *E. coli* K12 was diluted 1:100 into 50 mL LB medium and grown at 37 °C with shaking (200 rpm) for 16-18 h until reaching stationary phase. Cells were harvested by centrifugation (6000 × g, 10 min, 4 °C), washed twice with PBS (2 × 50 mL), and resuspended in LB medium to an $OD_{600}$ of 8. For compound treatment, 1 mL aliquots of the bacterial suspension were treated with 10 µL DMSO or compounds with desired concentrations for 1 h at 37 °C with shaking (200 rpm). The cells were subsequently collected by centrifugation (6000 × g, 10 min, 4 °C), washed with PBS (2 × 400 µL), and resuspended in 200 µL SDS solution (0.5% in PBS). Cell lysis was performed by sonication at 70% intensity for 30 s. The lysates were clarified by centrifugation (21,000 × g, 30 min, RT), and protein concentrations were determined using a BCA assay (Roti®Quant, Roth). Samples were adjusted to 1 µg/µL protein, and 44 µL aliquots were subjected to CuAAC labeling by addition of rhodamine azide (1 µL, 5 mM in DMSO), TBTA (3 µL, 1.667 mM in *t*-BuOH/DMSO, 4:1), TCEP (1 µL, 52 mM in $H_2O$), and $CuSO_4$ (1 µL, 50 mM in $H_2O$), followed by incubation at RT for 1 h.

The reaction was quenched with 50 µL 2× Laemmli buffer (63 mM Tris-HCl, 10% glycerol, 139 mM SDS, 0.0025% bromophenol blue, 5% 2-mercaptoethanol). Click reagents were removed by acetone precipitation using 400 µL acetone at -20 °C overnight. After acetone removal, protein pellets were resuspended in Laemmli buffer and heated at 95 °C for 10 min. Samples were separated by SDS-PAGE (12.5% acrylamide, 20 µL per lane). Roti®Mark Standard (Carl Roth) and BenchMark™ Fluorescent Protein Standard (Thermo Fisher) were used as protein size references. Fluorescence signals were recorded on a Fujifilm LAS-400 imaging system equipped with a Fujinon VRF43LMD3 lens and a 575DF20 filter. Total protein loading was assessed by Coomassie Brilliant Blue staining.

2.4. Preparative Labeling with **SCL-alkyne** and **STP-alkyne**[7],[8]

*In situ* labeling and lysis

An overnight culture of *E. coli* K12 was diluted 1:100 into 50 mL LB medium and grown at 37 °C with shaking (200 rpm) for 16-18 h until reaching stationary phase. Cells were harvested by centrifugation (6,000 × g, 10 min, 4 °C), washed twice with PBS (2 × 50 mL), and resuspended in LB medium to an $OD_{600}$ of 8. For compound treatment, 1 mL aliquots of the bacterial suspension were incubated with 10 µL DMSO or compounds with desired concentrations for 1 h at 37 °C with shaking (200 rpm). Cells were then collected by centrifugation (6,000 × g, 10 min, 4 °C), washed with PBS (2 × 400 µL), and resuspended in 200 µL SDS solution (0.5% in PBS). Cell lysis was performed by sonication at 70% intensity for 30 s. The lysates were clarified by centrifugation (21,000 × g, 30 min, RT), and protein concentrations were determined using a BCA assay (Roti®Quant, Carl Roth).

Click and enrichment

Samples were adjusted to a protein concentration of 2.23 µg/µL, and 45 µL of each sample, corresponding to 100 µg total protein, was transferred to a 96-well round-bottom plate (Greiner). For click reaction, a freshly prepared mixture of biotin azide (0.6 µL, 20 mM in DMSO), TBTA (2.5 µL, 1.667 mM in *t*-BuOH/DMSO, 4:1), TCEP (0.6 µL, 100 mM in $H_2O$), and $CuSO_4$ (1.2 µL, 50 mM in $H_2O$) was added to each sample (4.9 µL per sample). The reaction was incubated at 25 °C and 950 rpm for 90 min using a ThermoMixer®C (Eppendorf). Click reactions were quenched by adding 65 µL of 8 M urea containing 10 mM TCEP and 20 mM IAA, followed by incubation at 25 °C and 950 rpm for 15 min.

Subsequently, DTT (2 µL, 500 mM) and 2× hydrophobic/hydrophilic carboxylate-coated magnetic beads (10 µL, 1:1 mixture, Cytiva) were added. Proteins were precipitated onto the beads by addition of 175 µL ethanol. All subsequent steps were carried out on a Hamilton Microlab Prep system. After incubation for 5 min at RT and 500 rpm, beads were immobilized on a 96-well magnet plate (Alpaqua, MAGNUM FLX®), and the supernatant was removed. The beads were washed three times with 180 µL 80% ethanol and once with acetonitrile. Proteins were eluted twice with 75 µL 0.2% SDS in PBS at 40 °C and 800 rpm for 5 min. For enrichment, the eluates were incubated with streptavidin magnetic beads (50 µL, NEB) for 1 h at 25 °C and 800 rpm. After magnetic separation, beads were washed three times with 180 µL 0.1% NP-40 in PBS, twice with 6 M urea, and three times with 200 µL water.

Digestion and desalting

On-bead digestion was performed overnight at 37 °C and 900 rpm in 100 µL 50 mM TEAB containing trypsin (2 µL, 0.5 µg/µL, Promega). Peptides were recovered by washing the beads twice with 75 µL of 3% formic acid and desalted using in-house StageTips packed with SDB-RPS material (Empore™). The tips were equilibrated with 1% TFA in isopropanol before sample loading. Samples were then washed once with the same buffer and once with 0.2% TFA in water. Peptides were eluted with 1% ammonia in 80% acetonitrile, dried at 40 °C using a Concentrator plus (Eppendorf), and reconstituted in 30 µL 1% formic acid. For LC-MS/MS analysis, 3 µL of each sample was injected onto a timsTOF Pro mass spectrometer (Bruker) operated in DIA mode. All experiments were performed with four replicates.

2.5. Competitive Labeling with **Mitorubrin**, **Mitorubrinol** and **Mitorubrinol Acetate**

The experiment was performed according to the procedure described in the above preparative Labeling section. For competition experiments, 1 mL aliquots were first incubated with the parent compounds (**mitorubrin**, **mitorubrinol** and **mitorubrinol acetate**) in DMSO, or DMSO alone as control, for 1 hour at 37 °C with shaking at 200 rpm. The **SCL-alkyne** was then added, and incubation was continued for another 1 hour under the same conditions.

2.6. LC-MS/MS Measurements on TimsTOF Pro and Proteomic Data Analysis

Measurement parameters on timsTOF Pro

Peptides were analyzed using an UltiMate 3000 nanoLC system (Dionex) coupled to a timsTOF Pro mass spectrometer (Bruker) through a CaptiveSpray nano-electrospray ion source and a Sonation column oven. Samples were loaded onto a trap column (Acclaim PepMap 100 C18, 75 µm × 2 cm, 3 µm, Thermo Fisher Scientific) and desalted with 0.1% formic acid in water for 7 min at 5 µL/min. Peptides were then separated on an Aurora C18 analytical column (25 cm × 75 µm, 1.7 µm, IonOpticks) using a multi-step gradient at a flow rate of 400 nL/min. The gradient was set as follows: 5-17% solvent B over 36 min, 17-25% over 18 min, and 25-37% over 6 min, followed by washing at 95% B for 10 min before column re-equilibration. Mobile phase A was 0.1%

(v/v) formic acid in water, and mobile phase B was 0.1% (v/v) formic acid in acetonitrile. Mass spectrometric analysis was carried out in DIA-PASEF mode. The dual TIMS device was operated with accumulation and ramp times both set to 100 ms. MS1 scans covered an ion mobility range of 0.60-1.60 V·s/cm². For fragmentation, precursor ions were selected within a mass range of 400-1201 m/z and an ion mobility range of 0.60-1.43 V·s/cm². Each DIA-PASEF scan used two ion mobility isolation windows with a width of 26 m/z. In total, 32 isolation windows with 1 m/z overlap covered the full mass range, giving 16 dia-PASEF scans per MS1 scan and an overall cycle time of approximately 1.8 s. Collision energy was applied as a linear ramp from 59 eV at $1/K_0$ = 1.3 V·s/cm² to 20 eV at $1/K_0$ = 0.85 V·s/cm². TIMS calibration was performed with three Agilent ESI-L tuning mix ions (m/z 622, 922, and 1222), introduced through the CaptiveSpray source inlet to determine reduced ion mobility values ($1/K_0$).

Proteomic data analysis

Raw DIA data were processed with DIA-NN v1.8.1 in library-free mode.[9] An *in silico* spectral library was generated from the UniProt reference proteome of *E. coli* K12 (UP000000625, retrieved on 04.05.2024) using Trypsin/P digestion with up to two missed cleavages, together with deep-learning-based prediction of fragment spectra, retention times and ion mobilities. *N*-terminal methionine excision and cysteine carbamidomethylation were enabled, while variable modifications were not permitted. Peptides were restricted to 7-30 amino acids, precursor charge states of 2-4, precursor m/z values of 300-1800 and fragment ion m/z values of 200-1800. DIA-NN was set to generate a spectral library, quantitative matrices and a PDF report. Both “Precursor FDR (%)” and “Log level” were set to 1. In the “Algorithm” section, “Mass accuracy”, “MS1 accuracy” and “Scan window” were each set to 0. The options “Use isotopologues”, “MBR”, “Heuristic protein inference” and “No shared spectra” were selected. “Protein inference” was set to Genes, and the “Neural network classifier” was used in single-pass mode. For quantification, “Quantification strategy” was set to Robust LC (high precision), with “Cross-run normalization” set to RT-dependent. “Library generation” was set to smart profiling, and “Speed and RAM usage” was set to optimal results.

Volcano plot visualization

Following DIA-NN processing, LFQ intensity values were imported into Perseus 2.0.10.0[10] and $log_2$-transformed. Protein groups were retained, if three valid values were present in at least one group. Missing values were imputed from a normal distribution using the default parameters, with a width of 0.3 and a downshift of 1.8 applied to the total matrix. Statistical analysis was performed using a two-sample

Student's t-test with permutation-based multiple testing correction at an FDR of 0.05. Processed data tables were exported from Perseus, and final data visualization was performed in Python.

#### 2.7. isoDTB Experiments for Amino Acid Selectivity and Lysine-Residue Quantification [11]

*In situ* labeling and lysis

An overnight culture of *E. coli* K12 was diluted 1:100 into 50 mL LB medium and grown at 37 °C with shaking (200 rpm) for 16-18 h until reaching stationary phase. Cells were harvested by centrifugation (6000 × g, 10 min, 4 °C), washed twice with PBS (2 × 50 mL), and resuspended in LB medium to an $OD_{600}$ of 10. For each condition, two 300 µL aliquots of bacterial suspension, corresponding to parallel samples for heavy and light isoDTB tagging, were incubated with 3 µL of DMSO or compounds at the desired concentrations for 1 h at 37 °C with shaking at 200 rpm. Cells were then collected by centrifugation (6,000 × g, 10 min, 4 °C), washed with PBS (2 × 400 µL), and resuspended in 400 µL SDS solution (0.5% in PBS). Cell lysis was performed by sonication at 70% intensity for 30 s. The lysates were clarified by centrifugation (21,000 × g, 30 min, RT), and protein concentrations were determined using a BCA assay (Roti®Quant, Carl Roth).

Click with isoDTB tags and digestion

Samples were adjusted to a protein concentration of 1 µg/µL, and 200 µL of each sample, corresponding to 200 µg total protein, was transferred to 2 mL LoBind tubes (Eppendorf). For click reaction with isoDTB tags, a freshly prepared mixture of isoDTB heavy or light tags (4 µL, 50 mM in DMSO), TBTA (12 µL, 1.667 mM in *t*-BuOH/DMSO, 4:1), TCEP (4 µL, 50 mM in $H_2O$), $CuSO_4$ (4 µL, 50 mM in $H_2O$), and SDS solution (4 µL, 10% in PBS) was added to the corresponding heavy or light samples (28 µL per sample). The reaction was incubated at 25 °C and 300 rpm for 1 h. DNase I recombinant solution (1 µL, Roche) was then added to each sample, followed by incubation at 37 °C and 300 rpm for 30 min. The corresponding heavy and light samples were then added together. Subsequently, hydrophobic/hydrophilic carboxylate-coated magnetic beads (80 µL, 1:1 mixture, Cytiva) were added and the samples were incubated for 5 min. Proteins were precipitated onto the beads by addition of 1 mL ethanol. After incubation for 5 min at RT and 1,000 rpm, beads were immobilized on a magnetic rack (Magnetight™ Separation Stand, Novagen), and the supernatant was removed. The beads were washed three times with 800 µL of 80% ethanol and resuspended in 400 µL of 2 M urea containing 0.5% SDS in PBS. DTT (10 µL, 200 mM in water) was added, followed by incubation at 65 °C and 1,000 rpm for 15 min. IAA (10 µL, 400 mM in water) was then added, and the samples were incubated at 37 °C and 1,000 rpm for 30 min. Proteins were then again precipitated onto the beads by addition of 800 µL ethanol. The beads were washed three times with 800 µL of 80% ethanol and residual ethanol was removed. Finally, the beads were resuspended with 400 µL of 2M urea in PBS, trypsin (16 µL, 0.5 µg/µL, Promega) was added, and on-bead digestion was performed overnight at 37 °C and 300 rpm.

Enrichment and desalting

After digestion, samples were placed on the magnetic rack, and the supernatant was transferred to fresh 2 mL LoBind tubes (Eppendorf). Residual peptides were recovered from the beads by two additional elution steps with 100 µL 2% DMSO in water. For each elution, samples were incubated for 30 min at 37 °C and 1,000 rpm, placed on the magnetic rack, and the supernatant was combined with the previous peptide solution. For enrichment, freshly washed streptavidin agarose resin (100 µL resin in 1,200 µL PBS per sample, Thermo Fisher Scientific) was added, followed by incubation for 1 h at RT under rotation. Streptavidin resin was collected by centrifugation at 1,000 × g for 3 min, and the supernatant was removed. The beads were resuspended in 600 µL PBS and transferred to centrifuge columns (Fisher Scientific). The beads were washed by gravity flow twice with 600 µL PBS, three times with 600 µL water, and three times with 600 µL 50% acetonitrile in water. The columns were placed into fresh 1.5 mL LoBind tubes. Peptides were eluted by centrifugation at 3000 × g for 3 min with 250 µL 50% acetonitrile containing 0.1% TFA, followed by two additional elutions with 100 µL of the same solution. Peptides were dried at 45 °C using a Concentrator plus (Eppendorf) and reconstituted in 15 µL 0.1% TFA in water. For LC-MS/MS analysis, 7 µL of each sample was injected onto a timsTOF Pro mass spectrometer (Bruker) operated in DDA mode. All experiments were performed in four replicates.

2.8. Lysine-Residue Reactivity Profiling

An overnight culture of *E. coli* K12 was diluted 1:100 into 50 mL LB medium and grown to stationary phase for 16-18 h at 37 °C with shaking at 200 rpm. Cells were harvested by centrifugation (6,000 × g, 10 min, 4 °C), washed twice with PBS (2 × 50 mL), and resuspended in LB medium to an $OD_{600}$ of 10. For high-concentration labeling, 300 µL aliquots of the bacterial suspension were treated with **SCL-alkyne** at a final concentration of 100 µM for 1 h at 37 °C and 200 rpm. For low-concentration labeling, aliquots were treated with **SCL-alkyne** at a final concentration of 10 µM under the same conditions. In control experiments, both aliquots were labeled with 100 µM **SCL-alkyne**. Samples were then processed according to the isoDTB workflow described above.

2.9. LC-MS/MS Analysis of isoDTB Samples on TimsTOF Pro

The MS-measurement was performed with the same hardware as described in the previous section. 7 µL of each peptide sample were injected onto a timsTOF mass spectrometer (Bruker) coupled to an UltiMate 3000 nanoLC system (Dionex). Peptides were separated using solvent A, 0.1% formic acid in water, and solvent B, 0.1% formic acid in acetonitrile, with a flow rate of 400 nL/min. The LC gradient was run over 90 min, starting with 5% B for 7 min, followed by an increase to 40% B over 60 min, to 60% B over 5 min, and to 90% B within 0.1 min. The column was washed with 90% B for 9.9 min, returned to 5% B within 0.1 min, and re-equilibrated with 5% B for 7.9 min.

The mass spectrometer was operated in data-dependent PASEF mode. The dual TIMS analyzer was operated with equal accumulation and ramp times of 100 ms, and the $1/K_0$ ion mobility range was set to 0.85-1.40 V·s $cm^{-2}$. The capillary voltage of the CaptiveSpray source was set to 1500 V. Ten PASEF scans were acquired per TopN acquisition cycle, resulting in a total cycle time of 1.17 s. The mass range was set to m/z 100-1700. The precursor intensity threshold and target intensity were set to 1750 and 14,500 arbitrary units, respectively, and dynamic exclusion was applied for 0.4 min. The quadrupole isolation width was set to 2 Th for precursors below m/z 700 and to 3 Th for precursors above m/z 800. Collision energy was ramped linearly as a function of ion mobility, from 59 eV at $1/K_0$ = 1.6 V·s $cm^{-2}$ to 20 eV at $1/K_0$ = 0.6 V·s $cm^{-2}$. TIMS elution voltages were linearly calibrated using three ESI-L Tuning Mix ions (Agilent; m/z 622, 922, and 1222) spiked onto the CaptiveSpray source inlet filter to obtain calibrated $1/K_0$ values.

2.10. isoDTB Samples Data Analysis[12]

General setup of analysis software

The data were analyzed using FragPipe version 20.0. Database searches were performed against the UniProt reference proteome of *E. coli* K12 (UP000000625; retrieved on 04.05.2024), combined with reversed sequences. Further data processing was performed using the FragPipe-integrated tools MSFragger version 3.8[13], Philosopher version 5.0.0[14], IonQuant version 1.9.8[15], and Python version 3.9.13.

Analysis of the mass of modifications with FragPipe (open search)

The open search was configured with a precursor mass tolerance of -150 to 1,000 Da and an initial fragment mass tolerance of 20 ppm. Calibration and Optimization was set to “Mass calibration, parameter optimization”, and Isotope Error was set to “0”. The enzyme was specified as trypsin, with cleavage after “KR” and not before “P”.

Cleavage was defined as enzymatic, missed cleavages were set to 2, and Clip N-term M was enabled. The peptide length range was 6 to 50, and the peptide mass range was 500 to 5,000 Da. Fixed modifications were set to 0.0 Da, while variable modifications were retained as specified in the workflow. Crystal-C was disabled. For PeptideProphet, the settings were "--nonparam --expectscore --decoyprobs --masswidth 1000.0 --clevel -2". PTMProphet was not used. ProteinProphet was configured with "--maxppmdiff 2000000". Generate report was enabled using "--sequential --prot 0.01 --mapmods". Label-free quantification and TMT-Integrator were disabled, whereas IonQuant was enabled with labeling-based quantification. The following PTM-Shepherd settings were used: smoothing factor "2", precursor tolerance "0.01 Da", prominence ratio "0.3", peak-picking width "0.002 Da", localization background "4", and annotation tolerance "0.01 Da". Custom mass shift annotation was enabled using a reduced UniMod-derived mass shift list (This list can be found as "reducedMasslist.txt" in the PRIDE upload for this project). For PTM-Shepherd spectral annotation, b and y ion types were enabled, and the maximum fragment charge was set to 2. Spectral library generation was disabled. The complete FragPipe workflow containing these parameters was deposited in the PRIDE upload as "open_final.workflow". For downstream data analysis, the "global.modsummary.tsv" file was used. Values reported as "default-ptmshepherd-dataset (PSMs)" were plotted against "Theoretical Mass Shift" for mass shifts between 400 and 1,000 Da. The paired unannotated mass shifts of 755.3166 and 761.3226 Da for **SCL-alkyne**, or 561.3014 and 567.3108 Da for **STP-alkyne**, were retrieved for downstream analysis. All replicates were analyzed as the same experiment.

Analysis of amino acid selectivity with FragPipe (offset search)

For the Mass Offset Search, the following settings were used in MSFragger: precursor mass tolerance -20 to 20 ppm, fragment mass tolerance 20 ppm, Calibration and Optimization "Mass calibration, parameter optimization", Isotope Error "0/1/2", enzyme name "trypsin", cut after "KR", but not before "P", cleavage "enzymatic", missed cleavages "2", Clip N-term M enabled, peptide length 6 to 50, peptide mass range 500 to 5000 Da, variable modification of 57.02146 Da on C with max. 3 occurrences, no fixed modifications, and mass offsets of 755.3166/761.3226 Da for **SCL-alkyne** and 561.3014/567.3108 Da for **STP-alkyne**. Crystal-C was disabled. PeptideProphet was run with "--nonparam --expectscore --decoyprobs --masswidth 1000.0 --clevel -2". PTMProphet was disabled. ProteinProphet was run with "--maxppmdiff 2000000". Reports were generated with "--sequential --mapmods --prot 0.01". MS1 quantification was performed with IonQuant using labeling-based quantification. The light and heavy labels were set to 755.3166/761.3226 Da or

561.3014/567.3108 Da on any amino acid, respectively. The m/z, retention-time and ion-mobility windows were set to 10 ppm, 0.4 min and 0.05, respectively. Re-quantification was enabled. Top N ions, minimum frequency, minimum ions, minimum experiments and minimum isotopes were set to 3, 0.5, 2, respectively. Run TMT-Integrator was disabled. PTM-Shepherd was run with a smoothing factor of 2, precursor tolerance of 20 ppm, prominence ratio of 0.3, peak-picking width of 20 ppm, a minimum peak count of 10 PSMs, maximum fragment charge of 2, fragment mass tolerance of 20 ppm and PSM-based normalization. Annotation tolerance was set to 0.01 Da, and a custom reduced mass-list file was used (This list can be found as “reducedMasslist.txt” in the PRIDE upload for this project). Localization was performed using b and y ions with a localization background of 4. Spectral library generation was disabled. The complete FragPipe workflow containing these parameters was deposited in the PRIDE upload as “offset_final.workflow”.

The downstream data processing was adapted from the workflow described by Zanon et al. Output files from the four replicates were processed separately. Entries were first filtered against the corresponding psm.tsv file to retain only PSMs passing 1% PSM- and protein-level FDR filtering. Localization information was taken from the best_locs column. Only entries containing a single lower-case letter in best_locs were retained, and entries with a delta score ≤ 1 were removed. For each retained entry, the UniProt accession was extracted from the protein column and linked to the corresponding full protein sequence from the FASTA file. Peptides were mapped to the full protein sequence, and only peptides occurring exactly once in the protein were used for site assignment. The modified residue position was calculated from the peptide position and the localized residue position. Modifications at the protein *N*-termini, including position 2 after *N*-terminal methionine clipping, and at the protein *C*-termini were assigned as terminal modifications. A site identifier was generated from the UniProt accession, residue number and either the modified amino acid one-letter code or the label “N-term”/“C-term” for terminal modifications. Within each experiment, multiple PSM-level entries assigned to the same site identifier were counted once, resulting in one unique entry per modified site. Site lists from the individual replicates were then combined, and only sites detected in at least two of four replicates were retained for the final analysis. The number and percentage of modified sites were calculated for each amino acid and for terminal modifications.

Lysine-residue quantification with FragPipe (closed search)

For the Closed Search, MSFragger was used with a precursor mass tolerance of -20 to 20 ppm and a fragment mass tolerance of 20 ppm. Calibration and Optimization was set to “Mass calibration, parameter optimization”, and the isotope error was set to “0/1/2”. Trypsin was selected as the search enzyme, with cleavage after “KR” but not before “P”. Enzymatic cleavage was used, two missed cleavages were allowed, and *N*-terminal methionine clipping was enabled. The peptide length range was set to 6-50 amino acids, and the peptide mass range was set to 500-5000 Da. Carbamidomethylation of cysteine was set as a fixed modification, whereas oxidation of methionine and protein *N*-terminal acetylation were included as variable modifications. The light and heavy modifications of 755.3166/761.3226 Da for **SCL-alkyne** and 561.3014/567.3108 Da for **STP-alkyne**, were set as variable modifications on lysine, with one occurrence allowed for each modification. No mass offsets were used. Crystal-C was disabled. PeptideProphet was run with “--decoyprobs -- ppm --accmass --nonparam --expectscore”. ProteinProphet was run with “--maxppmdiff 2000000”. Reports were generated with “--sequential --prot 0.01”. MS1 quantification was performed with IonQuant using labelling-based quantification. The light and heavy labels were assigned to lysine using the corresponding mass shifts. The m/z, retention-time and ion-mobility windows were set to 5 ppm, 0.1 min and 0.01, respectively. Re-quantification was enabled, whereas match-between-runs and normalization were disabled. TMT-Integrator, PTM-Shepherd and spectral library generation were disabled. The complete FragPipe workflow containing these parameters was deposited in the PRIDE upload as “closed_final.workflow”.

The downstream data processing was adapted from the workflow described by Zanon et al. The “ion_label_quant.tsv” files from the four replicates were processed separately. For each entry, the modified peptide sequence was taken from either the “Light Modified Peptide” or the “Heavy Modified Peptide”, depending on the higher “PeptideProphet Probability”, and probe-related modification masses were replaced by an *asterisk. Peptides were mapped to the corresponding full-length protein sequence, and only peptides occurring once in the respective protein were used for site assignment. Modified lysine positions were determined and used to generate site identifiers in the format UniProt accession_K_residue number. When multiple modified peptide sequences were assigned to the same identifier, the shortest modified peptide and corresponding peptide sequence were retained. For each identifier, the Log2 ratio HL was calculated as an intensity-weighted mean of the corresponding ion-level $\log_2$-transformed heavy-to-light ratios, using the summed light and heavy ion intensities as weights. The values were discarded when the standard deviation of the Log2 ratio HL values exceeded 1.41. Replicate data were then combined, and only identifiers quantified in at least two of four replicates were retained. If different modified peptide sequences were observed for the same identifier across replicates, the shortest modified peptide and corresponding peptide sequence were kept. The expected Log2 ratio HL for engaged lysine sites was 0, corresponding to a 1:1 heavy/light ratio. The preferred quantification window was set to $-1 < \log_2(R) < 1$.

2.11. Analysis of Functional Sites

UniProt annotations for “Active site”, “Binding site”, “DNA binding site”, and “Site” were retrieved for proteins containing engaged lysine sites identified in the closed search. Residues annotated in these categories were defined as functional-site residues. The position of each engaged lysine site was compared with the annotated residues in the primary sequence. Sites directly overlapping with annotated functional residues were assigned to “Matched functional sites”, whereas non-annotated sites within five residues of an annotated functional residue were assigned to “Nearby functional sites”.

2.12. Microscopic Motility Tracking Assay[16]

An overnight culture of *E. coli* K12 was diluted 1:100 into 10 mL LB medium and grown at 37 °C with shaking (200 rpm) until reaching an $OD_{600}$ of 0.9-1.0. 100 µL aliquots of the bacterial culture were mixed with 1 µL of **SCL-alkyne** (5 and 10 mM) or DMSO as a vehicle control, and incubated for 30 min at 37 °C with shaking at 500 rpm on an Eppendorf shaker. Subsequently, 10 µL of each sample was transferred onto a microscopy slide and covered with a LifterSlip™ coverslip (*EMS*, catalog no. E72186-36). Bacterial movement was recorded at 37 °C for 10 s (10 fps) using a Leica DMi8 inverted microscope equipped with a Leica DFC365 FX camera (Wetzlar, Germany) and a 40 × objective lens. For each slide, four different positions were tracked. The experiment was performed in three independent replicates.

Image processing and motility analysis were carried out using a custom Python script available at https://github.com/MIsselstein/E.Coli_Tracker.[16] The images were defined as binary images and combined into movies comprising 99 frames per 10 s recording. For bacterial cells tracking, the search range, corresponding to the maximum distance a detected feature was allowed to move between consecutive images, was set to 20. The memory parameter, which defines the maximum number of frames over which a feature may disappear and still be re-linked to the same bacterial cell upon reappearance, was set to 5. The magnification and pixel size were set to 40 and 6.45, respectively. Detected trajectories were exported both as movies and Excel files for subsequent analysis.

2.13. Chemotaxis Capillary Assay[17]

An overnight culture of *E. coli* K12 was diluted 1:100 into 20 mL LB medium to an $OD_{600}$ of 0.05 and grown at 37 °C with shaking (200 rpm) until reaching an $OD_{600}$ of 0.5. Cells were harvested and washed twice with PBS by centrifugation (2,000 × g, 10 min), resuspended in PBS, and adjusted to an $OD_{600}$ of 0.1. Then, 1 mL of the bacterial suspension was mixed with 10 µL of 10 mM **SCL-alkyne** or DMSO as a vehicle control. Aliquots of 200 µL were transferred to a 96-well plate and incubated for 1 h without shaking. The plate was sealed with three layers of Parafilm®M (Amcor). Flame-sealed 1 µL capillaries (Microcaps®, catalog no. #612-2470) prefilled with PBS or 100 mM glucose in PBS were inserted into the bacterial suspension. After incubation for 1 h at room temperature, the exterior of each capillary was wiped with Bacillol®AF (HARTMANN) to remove adherent bacteria. The liquid inside each capillary was then expelled into 400 µL PBS, and 50 µL of the resulting suspension was plated on LB agar. Plates were incubated at 37 °C for 15 h, and colony-forming units (CFU) were counted.

2.14. UV-Visible Absorption Spectra with or without $FeCl_2$

$FeCl_2$ solutions were freshly prepared in Milli-Q water at concentrations of 100 µM or 1 mM. **Mitorubrin** in DMSO (10 mM) was added to the $FeCl_2$ solutions to obtain a final **mitorubrin** concentration of 100 µM. **Mitorubrin** alone and $FeCl_2$ alone were included as controls. Samples were transferred to UV-transparent 96-well plates (UV-Star®, Greiner) with 200 µL per well. UV-visible absorption spectra were recorded from 230 to 500 nm at 25 °C using an Infinite M200 PRO microplate reader (Tecan Group).[18]

2.15. Determination of Minimal Inhibitory Concentration (MIC)

The minimal inhibitory concentration was determined using a broth microdilution assay. Assays were performed in LB medium or LB supplemented with 1 mM $FeCl_3$. To reduce evaporation, the outer wells of sterile flat-bottom 96-well plates (Greiner) were filled with 200 µL of LB medium. **Mitorubrin** (10 mM in DMSO) was diluted in LB medium, followed by two-fold serial dilution in LB containing 2% DMSO, with or without 1 mM $FeCl_3$. Bacterial overnight cultures were diluted 1:10,000 in LB medium with or without 1 mM $FeCl_3$, and 50 µL of the inoculum was added to each well, resulting in a final assay volume of 100 µL. The final DMSO concentration in each well was 1%. Sterile and growth controls were included on each plate. Plates were incubated at 37 °C and 200 rpm for 24 h. The MIC was defined as the lowest compound concentration at which no visible bacterial growth was observed. MIC values were determined in three technical replicates and confirmed in two independent biological replicates.

#### 3. Supplementary Figures

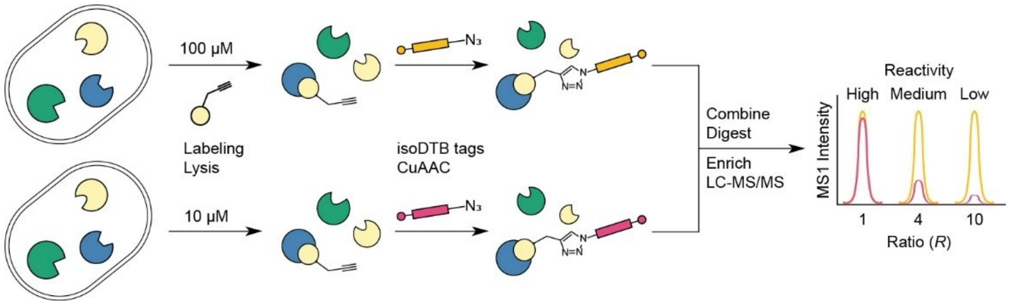

**Figure S1.** Workflow for measuring lysine reactivity using isoDTB tags. Classification of lysine reactivity based on isotopic $R_{10:1}$ ratios: high reactive ($R_{10:1}$ < 3), medium reactivity (3 < $R_{10:1}$ < 5), and low reactivity sites ($R_{10:1}$ > 5).

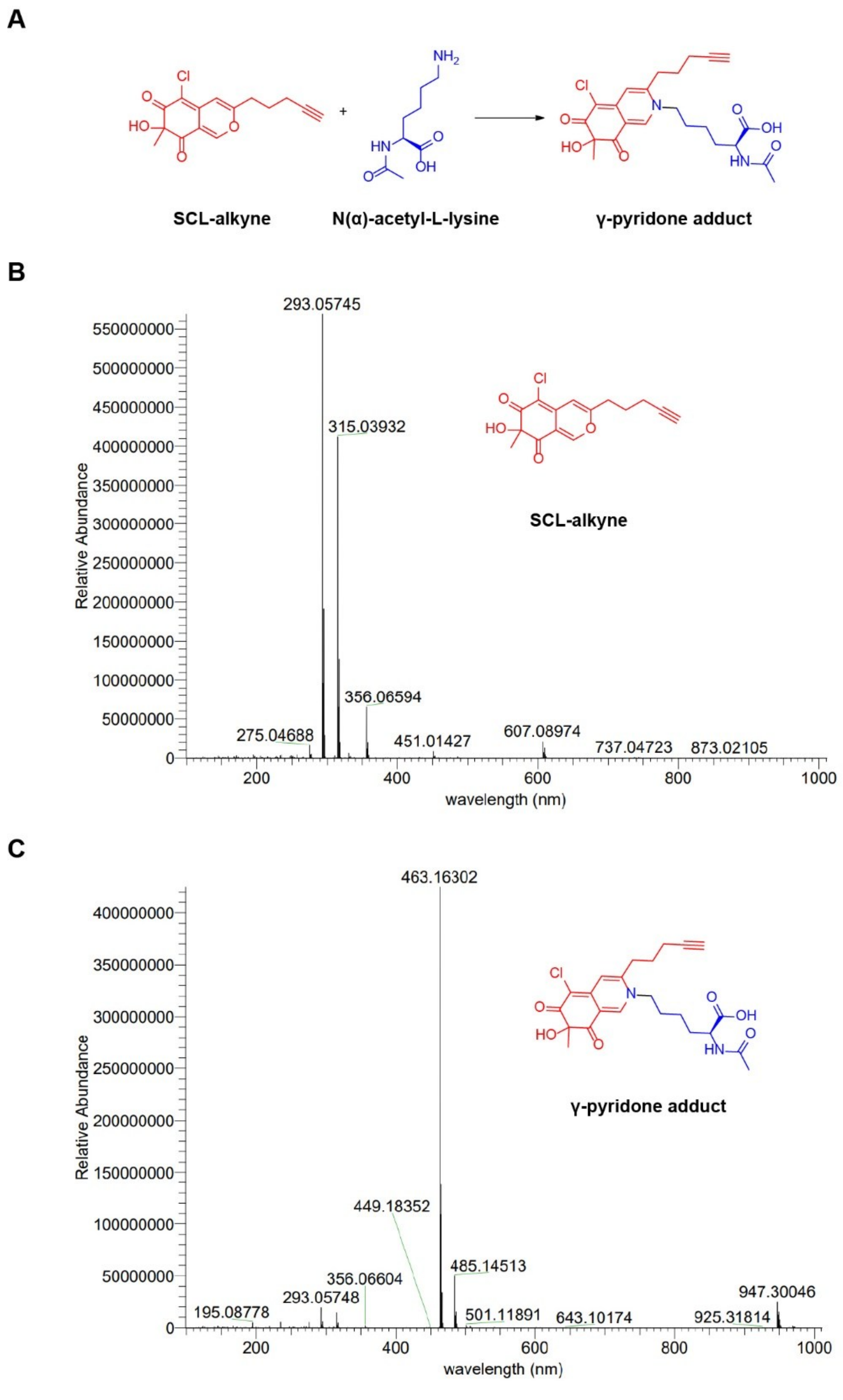

**Figure S2.** (A) Reaction of **SCL-alkyne** (2.0 mM) with *N*(α)-acetyl-L-lysine (1.0 mM) in phosphate buffer (100 mM, pH 8.0) at 37 °C. Formation of the corresponding γ-pyridone adduct was monitored by HPLC-MS. (B) Mass spectrum of **SCL-alkyne** before reaction, showing the protonated ion at m/z 293.0575. (C) Mass spectrum of the reaction mixture after 1 h, showing the γ-pyridone adduct ion at m/z 463.1630.

**A**

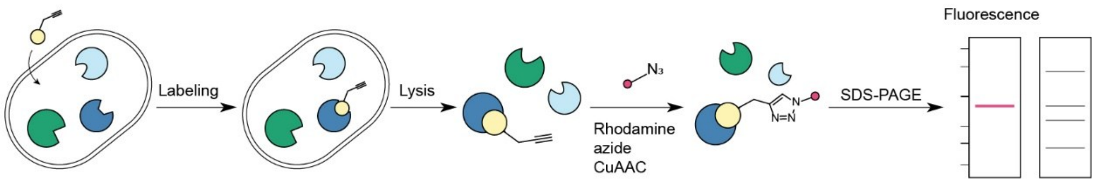

**B**

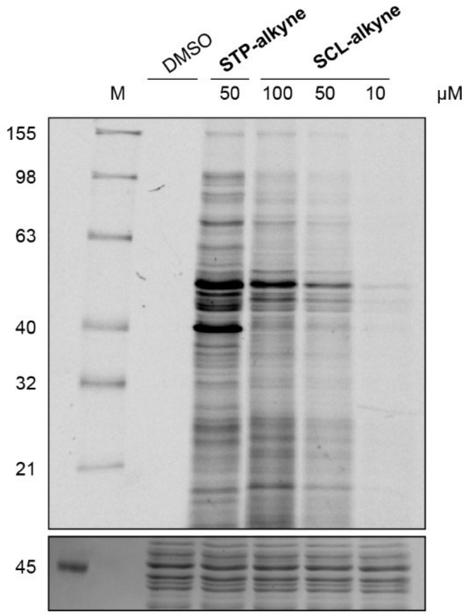

**Figure S3.** Workflow for protein target visualization. Live *E. coli* K12 cells were incubated with **SCL-alkyne**, **STP-alkyne** or DMSO as a control. Probe-labeled proteins were subsequently visualized by in-gel fluorescence after CuAAC with rhodamine azide. (B) Concentration-dependent labeling of proteins in intact cells was observed after 1 h incubation with **SCL-alkyne**, with 50 µM **STP-alkyne** included as a benchmark. Fluorescently labeled proteins were detected in-gel, and Coomassie staining was included as a loading control.

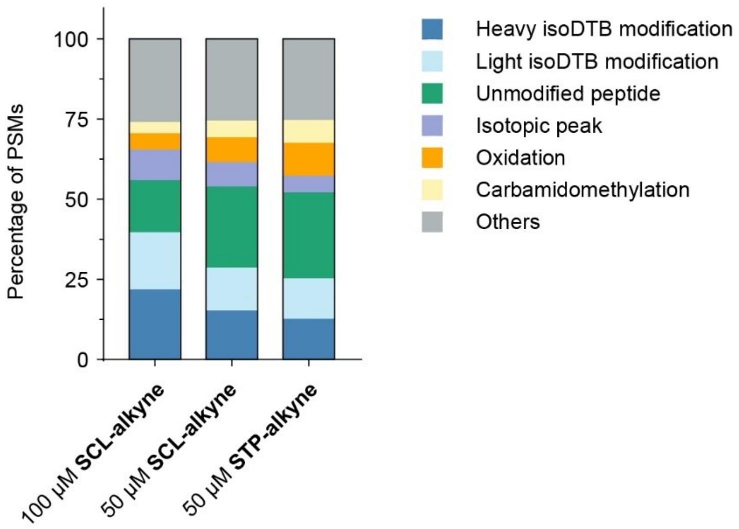

**Figure S4.** The plot shows the percentage of PSMs assigned to different modifications in the Open Search analysis. Major modifications are shown individually, and all modifications accounting for less than 3% were grouped as “Others”. Detailed percentage values are provided in Table S2.

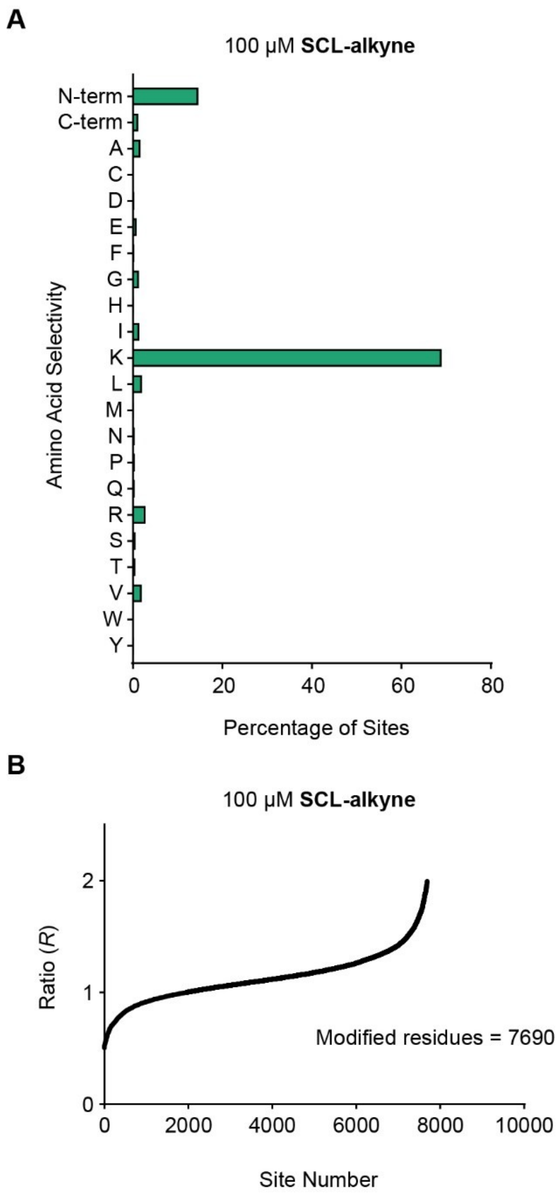

**Figure S5.** (A) The bar graph shows the distribution of modified amino acid residues, including *C*-terminal (C-term) and *N*-terminal (N-term) modifications, in *E. coli* K12 cells treated with 100 μM **SCL-alkyne**. (B) *R* values of lysine residues labeled by **SCL-alkyne** at 100 μM in *E. coli* K12 cells were determined from four technical replicates. Only lysine sites with *R* values between 0.5 and 2 were considered engaged sites. See Table S4 for the lysine-residue quantification data.

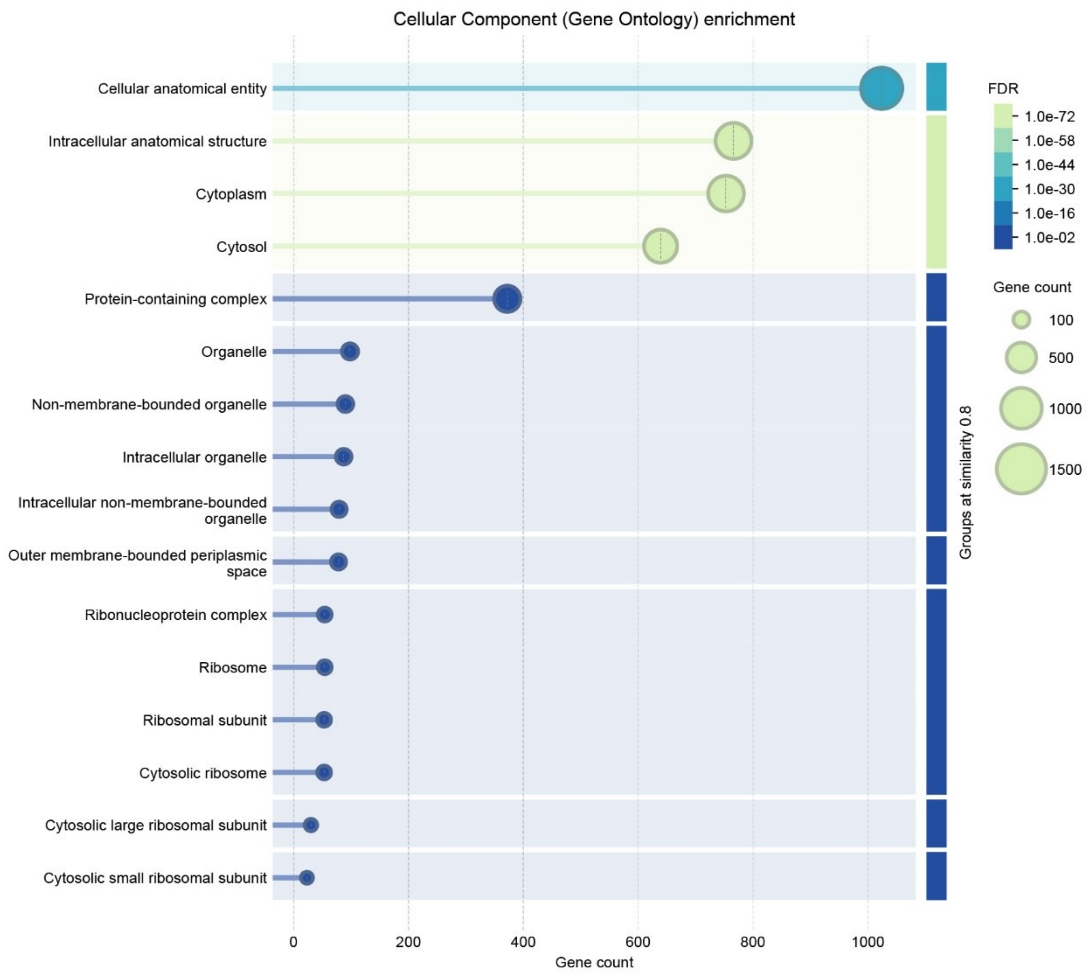

**Figure S6.** Gene ontology analysis of cellular component with 50 µM **SCL-alkyne** binding proteins from closed search in the above isoDTB workflow. This was analyzed via String with Group Similarity ≥ 0.8 and medium confidence ≥ 0.4.

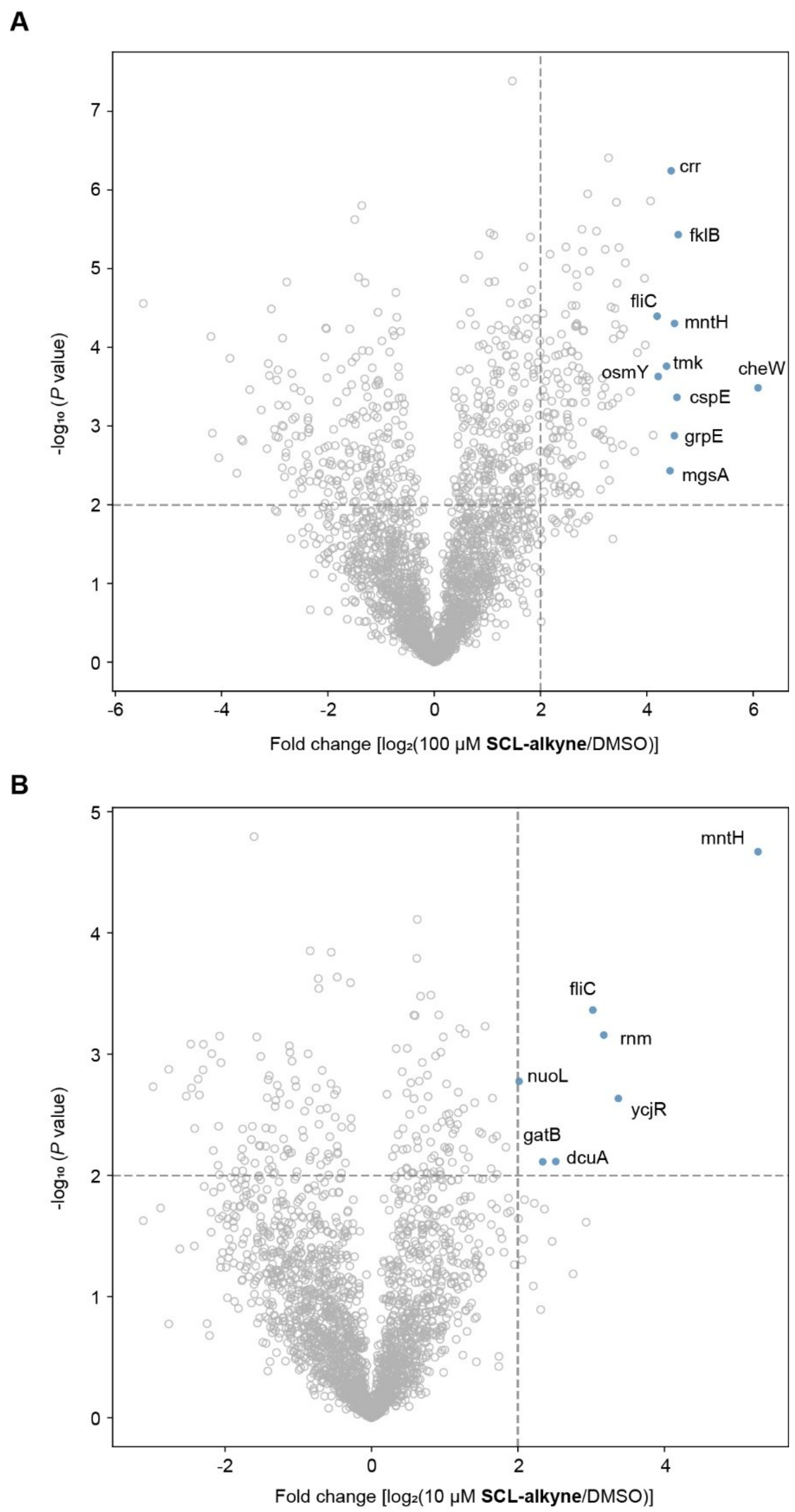

**Figure S7.** Volcano plots showing proteins enriched after treatment with (A) **SCL-alkyne** (100 µM, 1 h) or (B) **SCL-alkyne** (10 µM, 1 h). Thresholds were set at $\log_2$ fold change > 2.0 and p-value < 0.01 (two-tailed Student's t-test, n = 4 replicates per group). The top 10 most enriched proteins were annotated with their corresponding gene names. See Table S1 for the list of identified proteins.

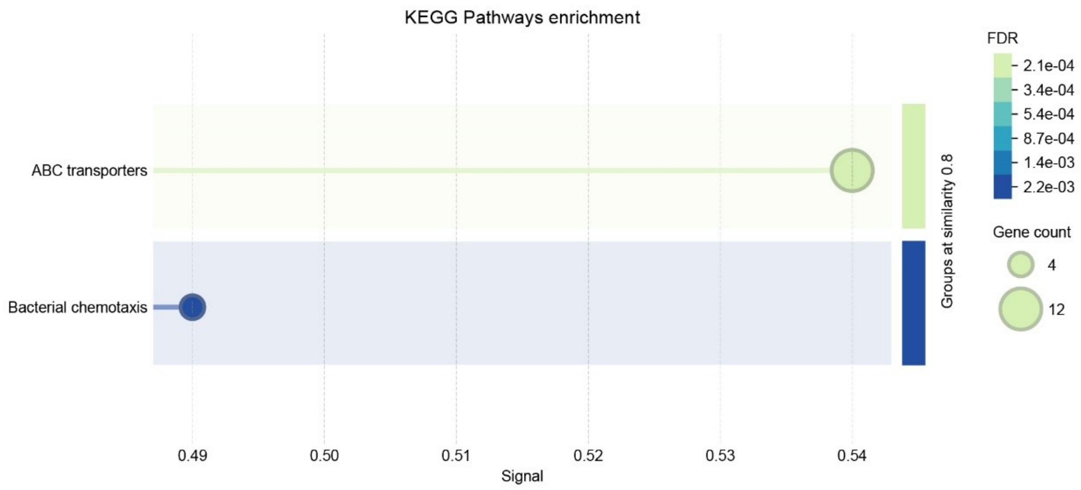

**Figure S8.** KEGG pathway enrichment analysis of proteins enriched by 50 μM and 100 μM **SCL-alkyne**. This was analyzed via String with Group Similarity ≥ 0.8 and medium confidence ≥ 0.4. Signal score reflects the overall enrichment intensity by balancing enrichment strength and the false discovery rate.

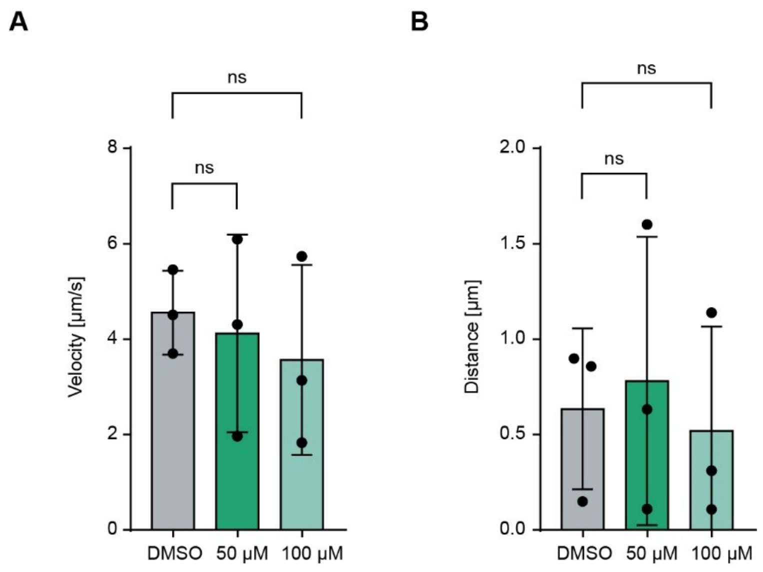

**Figure S9.** Comparison of swimming velocity (A) and distance (B) between untreated and **SCL-alkyne** treated (50 and 100 µM) *E. coli* K12 cells in motility tracking experiments. Data are presented as mean ± SD from three independent replicates (n = 3). Statistical significance was determined using one-way ANOVA. ns, not significant.

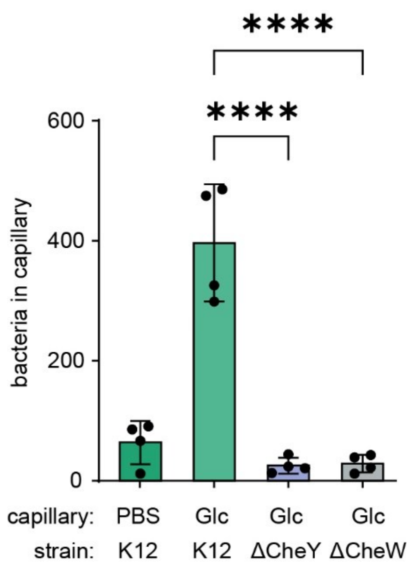

**Figure S10.** The number of bacteria was determined by counting colonies on agar plates. The strains used here were *E. coli* K12, *E. coli* BW2511 Δ*cheY* and Δ*cheW* mutant strains. Glc, glucose. Data are shown as mean ± SD from four biological replicates (n = 4). Statistical significance was determined using one-way ANOVA. (****p ≤ 0.0001).

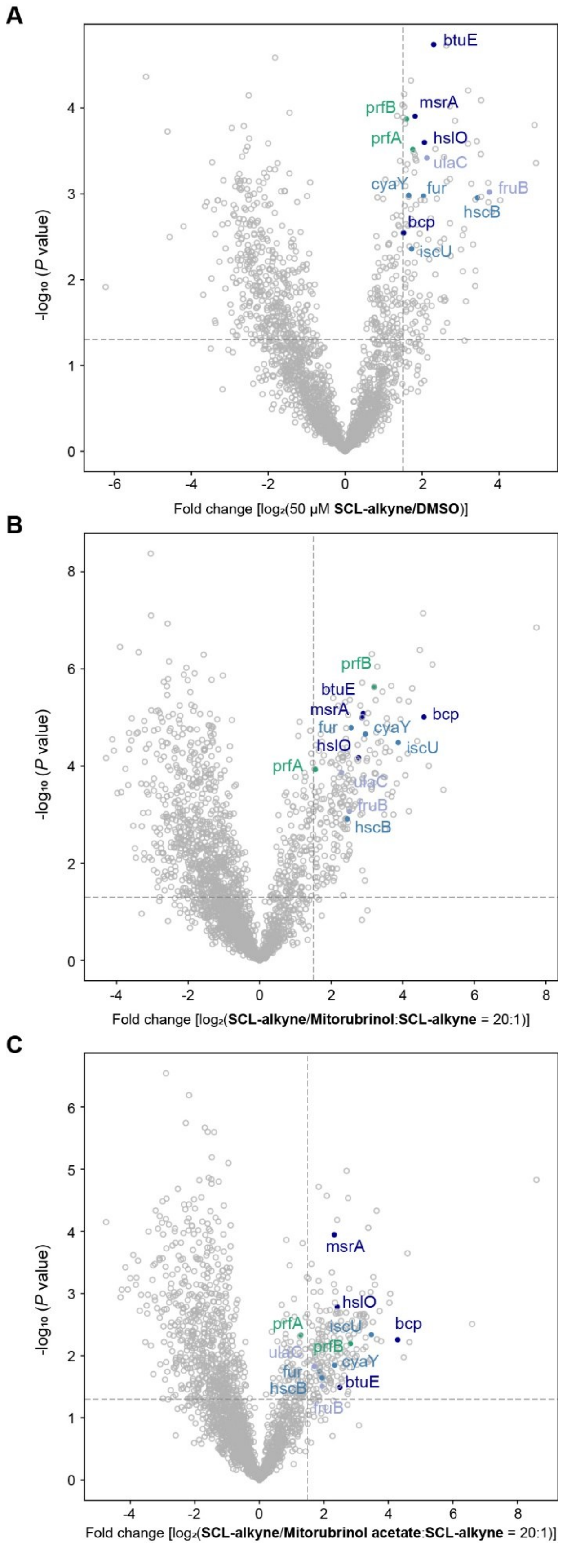

**Figure S11.** Volcano plot of competitive ABPP in intact *E. coli* K12 cells. The plot shows preparative labeling results of cells treated with **SCL-alkyne** and competition by mitorubrin natural products. (A) Cells treated with **SCL-alkyne** (50 µM, 1 h) compared with DMSO controls. (B and C) **SCL-alkyne** labeling alone (50 µM, 1 h) compared with **SCL-alkyne** labeling after preincubation with 1 mM **mitorubrinol** (B) or 1 mM **mitorubrinol acetate** (C). Thresholds were set at $\log_2$ fold change > 1.5 and p-value < 0.05 (two-tailed Student's t-test, n = 4 replicates per group). Proteins belonging to the clusters identified in the heat map were annotated with their gene names and color-coded according to biological process. Dark blue: oxidative stress; light blue: iron-sulfur cluster assembly; green: protein-containing complex disassembly; light purple: phosphorylation. See Table S6 for the list of identified proteins.

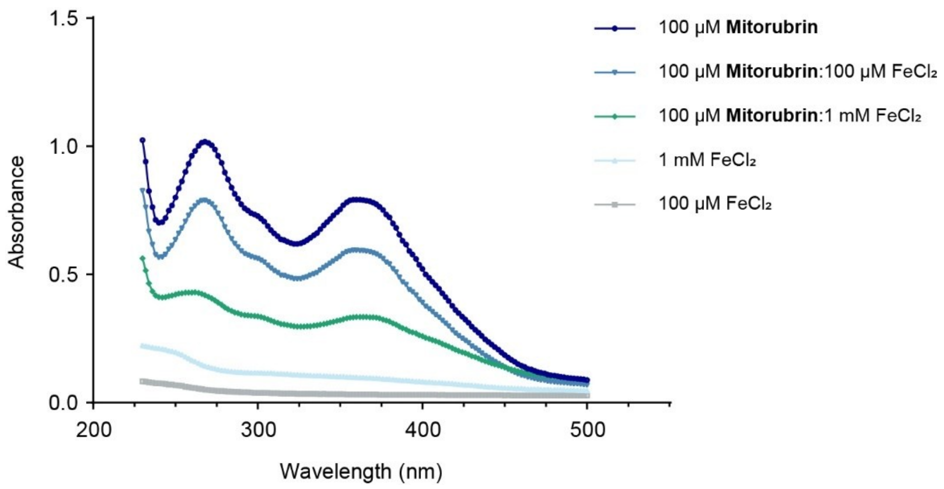

**Figure S12.** UV-visible absorption spectra of 100 μM **mitorubrin** alone and in mixtures with $FeCl_2$ at ratios of 1:1 and 1:10. $FeCl_2$ alone at 100 μM and 1 mM was included as background controls.

#### 4. Supplementary Tables

see separate Excel files

**Table S1.** Chemical proteomics dataset of **SCL-alkyne** and **STP-alkyne** in *E. coli* K12 cells including protein annotation, LFQ intensities, and statistical analysis.

**Table S2.** Mass of modification data for **SCL-alkyne** and **STP-alkyne**, determined by open search using the MSFragger-based FragPipe workflow.

**Table S3.** Amino acid selectivity data for **SCL-alkyne** and **STP-alkyne**, determined by offset search using the MSFragger-based FragPipe workflow.

**Table S4.** Lysine-residue quantification data for **SCL-alkyne** and **STP-alkyne**, determined by MSFragger closed search and IonQuant labeling-based quantification.

**Table S5.** Lysine-residue reactivity data for the experiments comparing 10 µM and 100 µM as well as 100 µM and 100 µM **SCL-alkyne** in *E. coli* K12 cells.

**Table S6.** Chemical proteomics data of **SCL-alkyne** labeling in competition with compounds **mitorubrin**, **mitorubrinol** and **mitorubrinol acetate** in *E. coli* K12 cells including protein annotation, LFQ intensities, and statistical analysis.

#### 5. NMR Spectra

(*E*)-4-((methoxyimino)methyl)-2-methyl-1,3-phenylene dibenzoate (**3**)

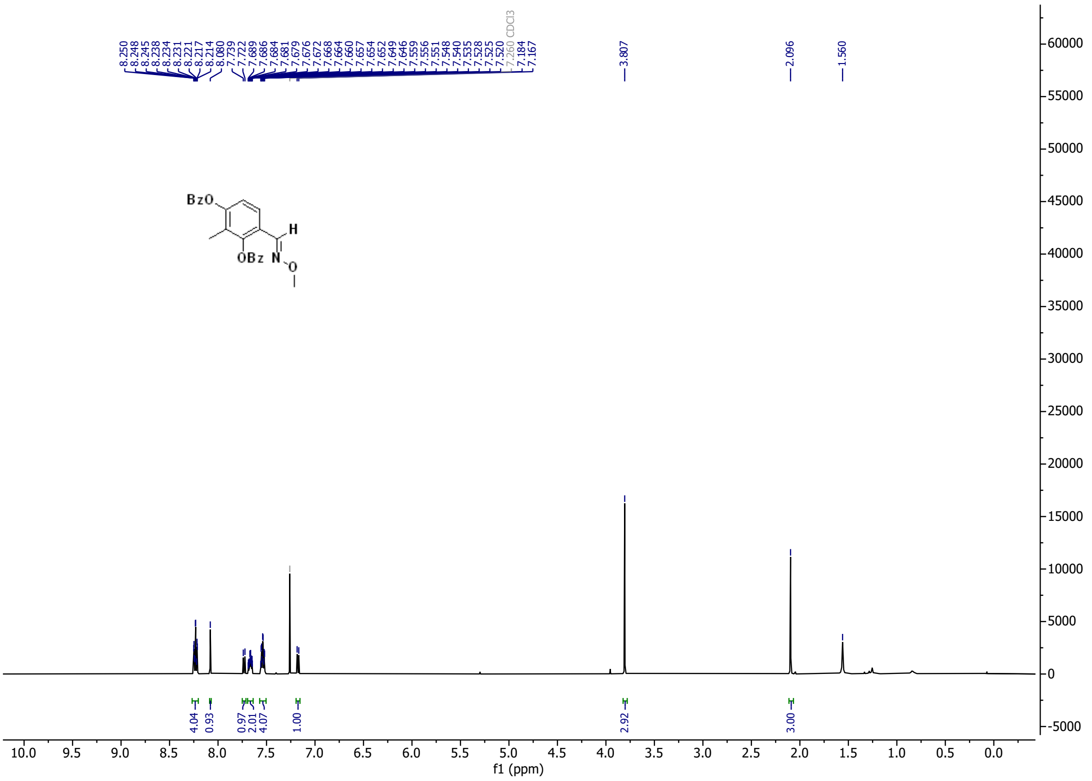

(*E*)-5-bromo-4-((methoxyimino)methyl)-2-methyl-1,3-phenylene dibenzoate (**4**)

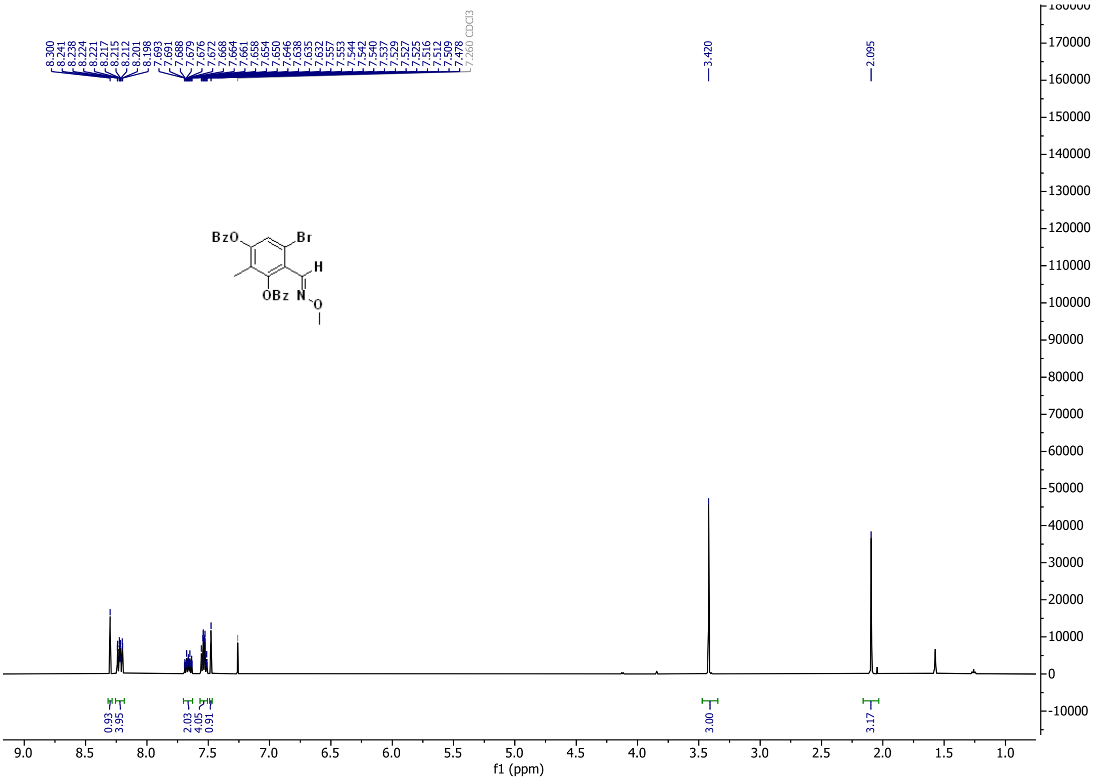

6-bromo-2,4-dihydroxy-3-methylbenzaldehyde (**6**)

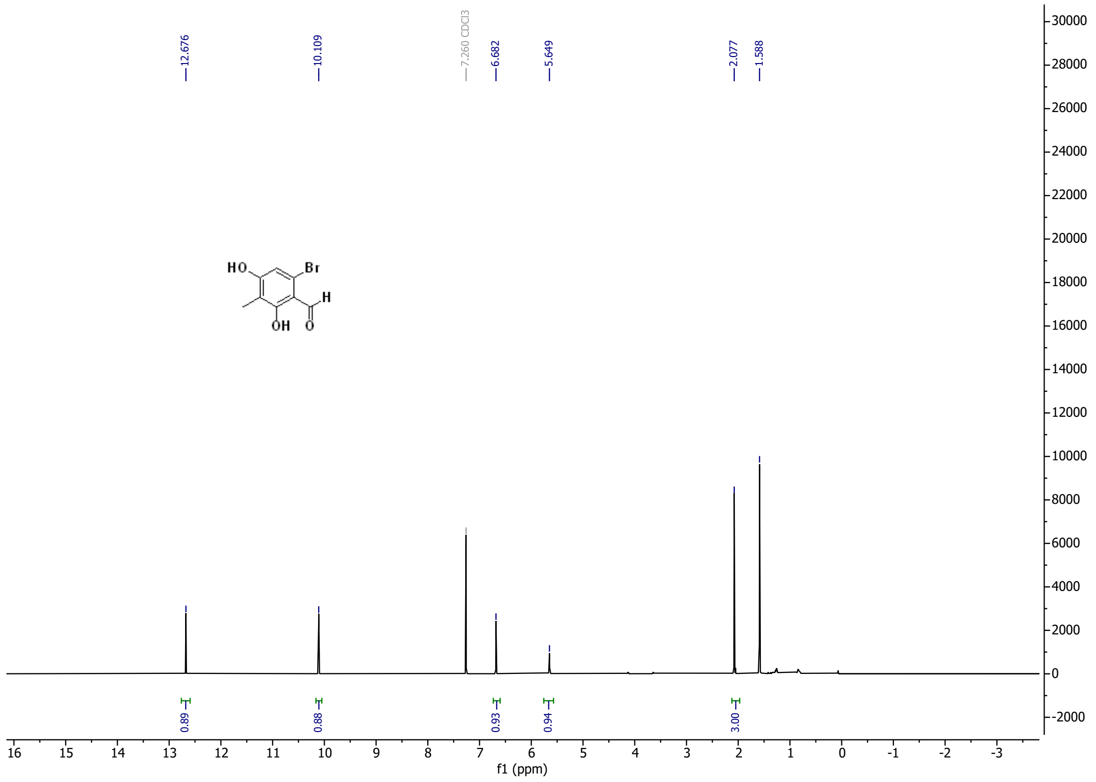

6-(hepta-1,6-diyn-1-yl)-2,4-dihydroxy-3-methylbenzaldehyde (**7**)

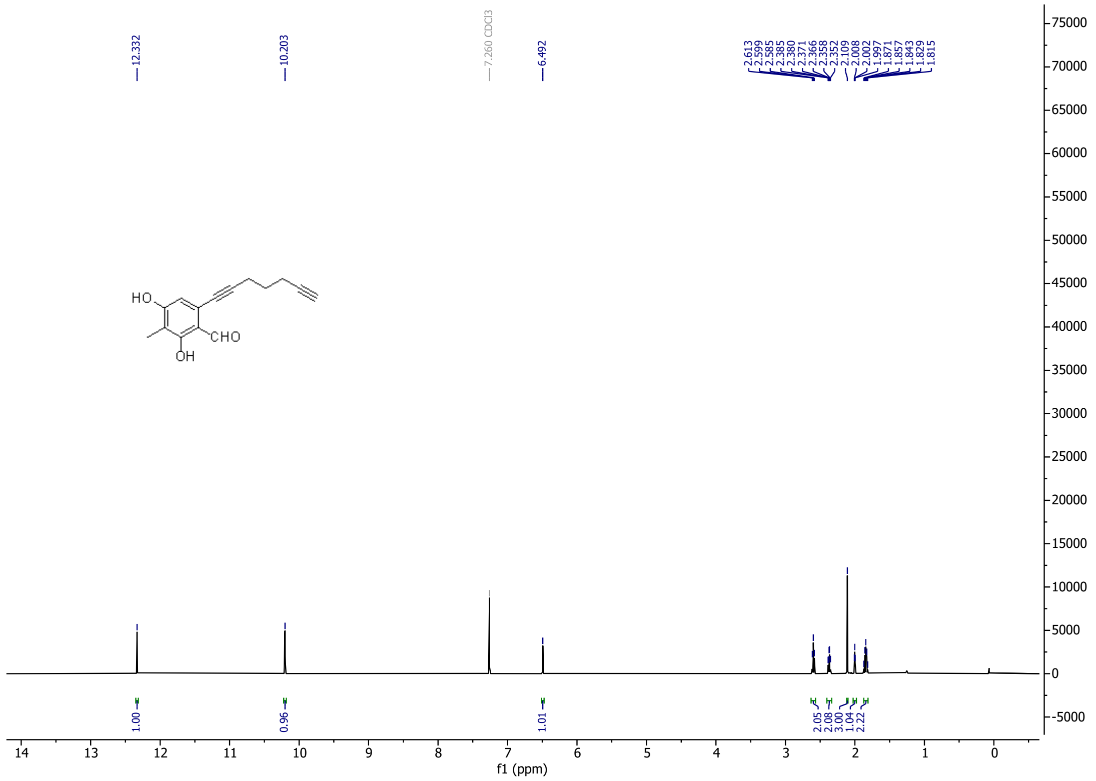

7-hydroxy-7-methyl-3-(pent-4-yn-1-yl)-6*H*-isochromene-6,8(7*H*)-dione (**8**)

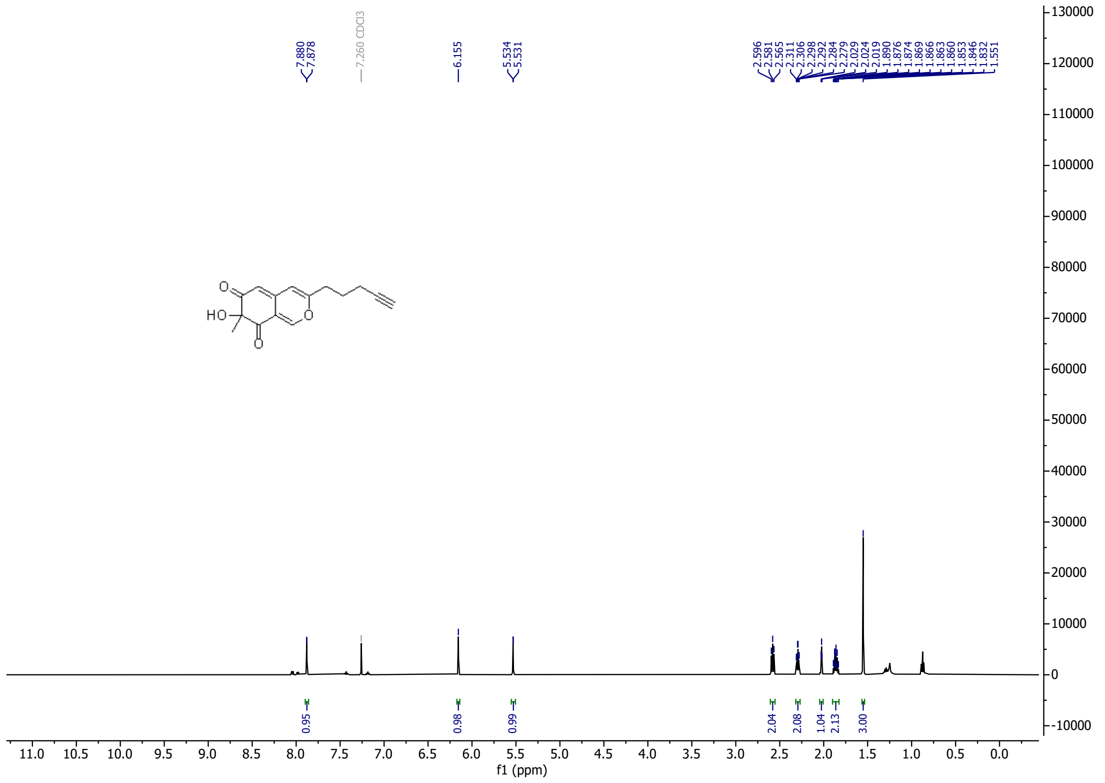

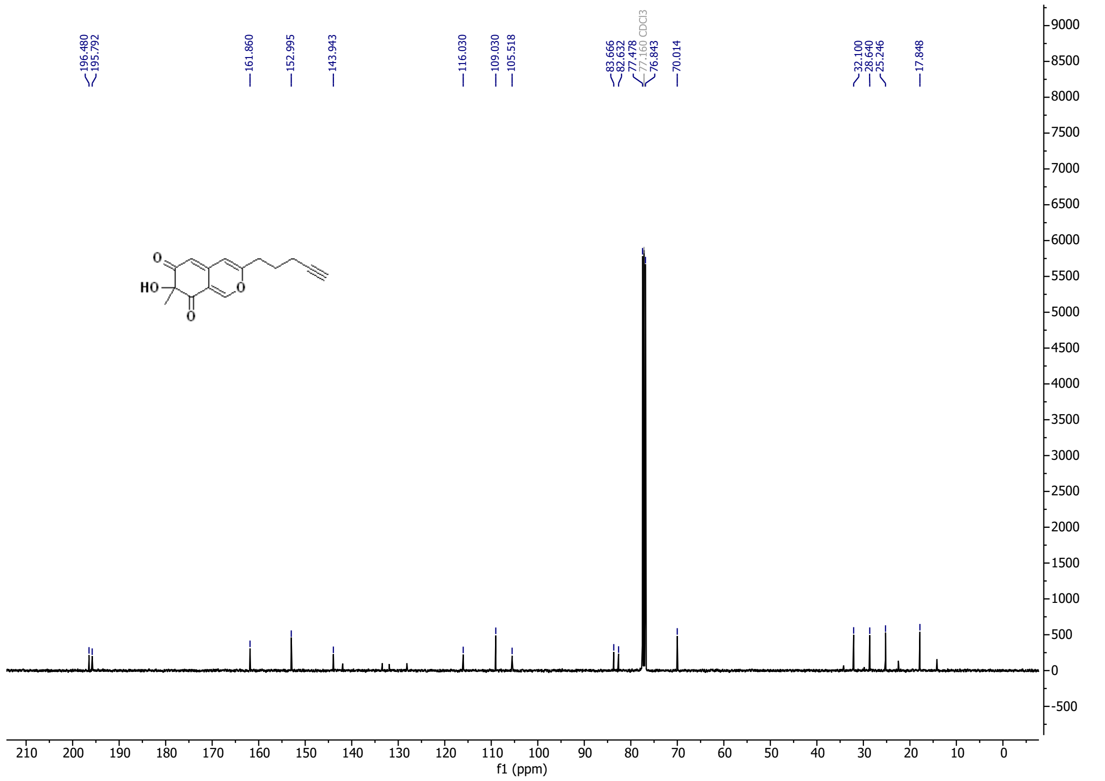

5-chloro-7-hydroxy-7-methyl-3-(pent-4-yn-1-yl)-6*H*-isochromene-6,8(7*H*)-dione (**SCL-alkyne**)

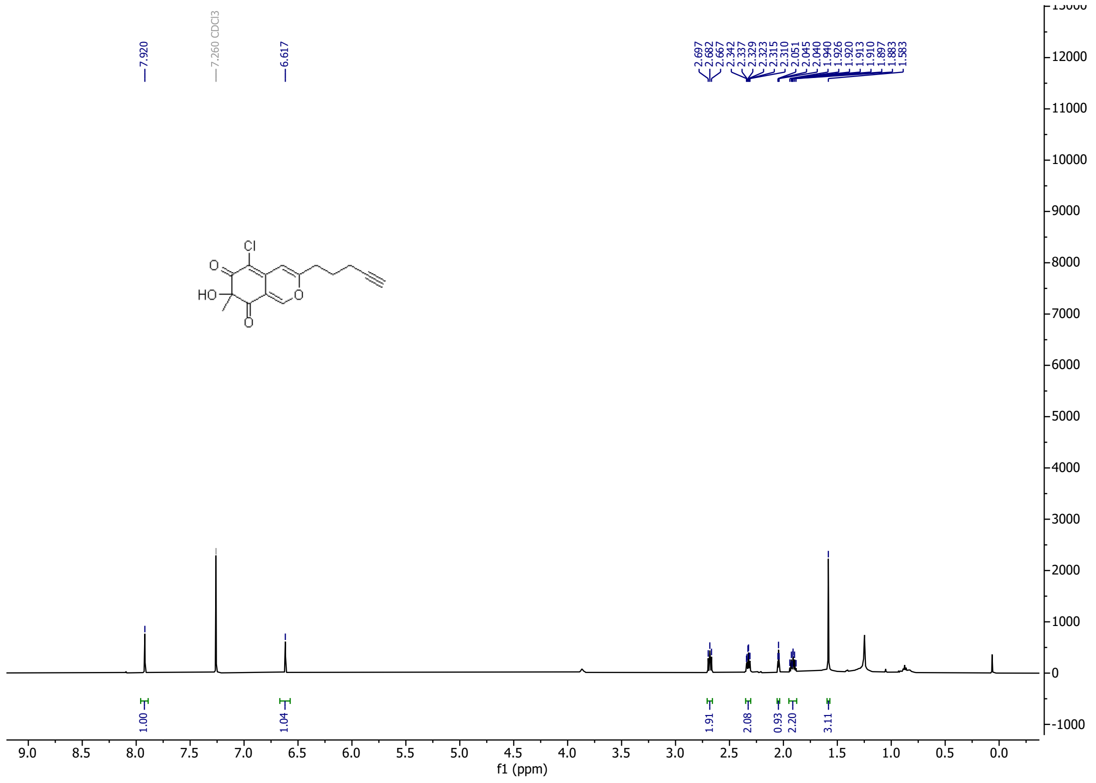

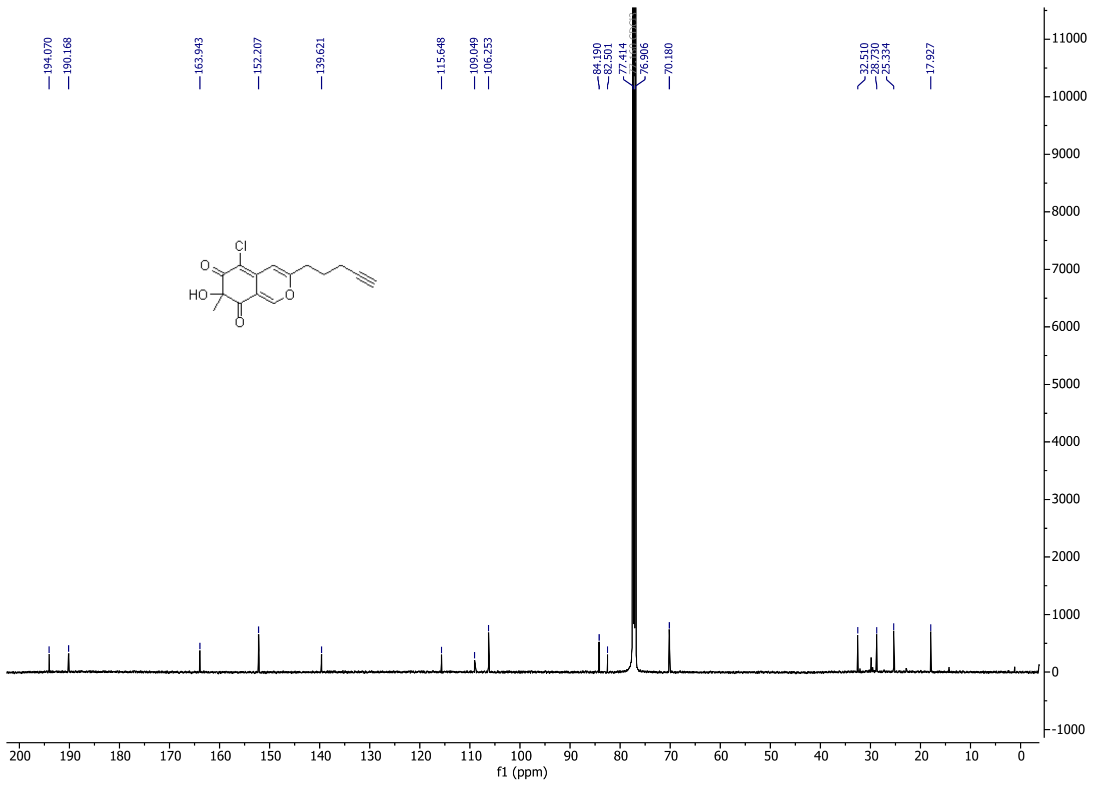
